# Repetition-related reductions in neural activity support improved behavior through increases in oscillatory power

**DOI:** 10.1101/2025.07.06.663291

**Authors:** Adrian W. Gilmore, Leonardo Claudino, Cassandra M. Levesque, Anna M. Agron, Peter J. Molfese, Vinai Roopchansingh, Michael D. Rugg, Stephen J. Gotts, Alex Martin

## Abstract

Repeatedly processing an object leads to subsequent behavioral improvements in its identification and reductions in associated neural activity, a form of neural processing efficiency that multiple theoretical models have attempted to explain. Using simultaneous fMRI-EEG in humans during object naming, we find that stimulus-driven oscillatory power from high (gamma) to low frequencies (theta) increases and resolves earlier with repetition in task-engaged frontal and occipitotemporal brain regions identified as showing repetition suppression in fMRI. Changes in gamma oscillatory power in these regions were correlated with behavioral priming across subjects and additionally with fMRI repetition suppression in left frontal cortex, providing multimodal support for a novel mixture of the previously proposed Synchrony and Facilitation models.

## INTRODUCTION

It is a foundational phenomenon in cognitive neuroscience that repeated experiences with a stimulus lead to behavioral improvements in the task being performed (repetition priming) while simultaneously leading to reductions in neural activity (repetition suppression; RS). This dual neural/behavioral phenomenon has been observed in humans^1,2^, non-human primates^3,4^, and rodents^5,6^. Across this range of species, repetition-related changes consistently appear to be stimulus-specific and have both short-term and long-term components^7^. The phenomenon is also notable in that it does not depend strongly on the hippocampus and other medial temporal lobe structures, as it is largely spared in amnesia^8–10^. Rather, it prominently involves the neocortical regions engaged by the task being performed. Left lateral prefrontal cortex appears to play a particularly critical role in mediating the relationship between neural and behavioral effects^11–14^, possibly through the mapping of behavioral responses to specific perceptual inputs^15,16^.

However, the co-occurrence of priming and RS poses a mechanistic puzzle. How might reducing neural activity support improvements in—rather than impairments of—the behavior in question? “Efficiency” is often invoked to describe the phenomenon, but it does not, by itself, imply any specific mechanism^17^. However, several theoretical models have been proposed to explain this coupling. The Facilitation model^18,19^ proposes that neural activity initiates and resolves more quickly for repeated stimuli, with rapid kinetics that are difficult to capture given the slow timecourse of BOLD fMRI responses when studied in humans. The Sharpening model^1,20^ holds that RS occurs primarily in cells that are poorly tuned to the repeated stimulus, thereby increasing neural selectivity despite an overall activity reduction. The Synchrony model^7,21^ holds that cells fire in a more synchronized manner with repetition, leading to larger amplitude population fluctuations that require fewer spikes for effective signal propagation.

Finally, the Predictive Coding model^22^ holds that the brain is composed of generative neural networks that attempt to minimize differences over repetitions between top-down expectations and bottom-up sensory evidence, with observed neural activity reflecting residual prediction error. While all models have some experimental support, none has yielded strong direct empirical evidence that ties their proposed mechanisms to the magnitude of both RS and repetition priming^23^.

In humans, one of the main limitations to progress has been a lack of neuroimaging methods that have both high spatial and temporal resolution with full brain coverage—both of which seem required to better understand the relation between behavioral improvements and reductions in neural activity. In the current study, we address this limitation by utilizing simultaneous fMRI-EEG while participants name pictures of common objects that are either novel (presented a single time) or repeated during the experiment. fMRI provides a spatially precise means of identifying brain regions that exhibit RS. The simultaneously acquired EEG data can then be source localized to the cortex at the anatomical locations identified by fMRI, providing an estimate of rapid electrophysiological neural activity at those locations that cannot be obtained by BOLD fMRI alone.

## RESULTS

### Repetition priming, fMRI repetition suppression, and scalp ERP repetition effects are observed in object naming

Forty participants overtly named 100 pictures of common objects (living things; inanimate objects) twice each, approximately one hour prior to brain imaging. During fMRI-EEG, participants then named the same 100 pictures once more (Repeat condition), randomly intermixed with a set of 100 new objects (Novel condition) that were matched in lexical properties and name frequency to the repeated set (Fig. 1A). Accuracy and response times were recorded for each trial. As in prior studies, repeated pictures were named faster than novel pictures [Repeat *M*(SD) = 853.1 (16.6) ms, Novel *M*(SD) = 952.3 (21.3) ms; *t*(39) = 11.46, *p < .*001, Cohen’s *d* = 1.81; Fig. 1B]. Accuracy did not differ significantly between conditions (*p = .*234). Also consistent with prior studies^23,24^, picture naming led to above-baseline activity in fMRI in occipitotemporal and frontal brain regions (Fig. 1C), with RS prominently observed for repeated pictures in left lateral frontal cortex, anterior cingulate cortex (extending into the pre-supplementary motor area; hereafter “dACC/pre-SMA”), and the left and right fusiform gyri (all *p*s *< .*001, FDR-corrected to *q* < .05; Fig. 1D). In the simultaneously acquired EEG, scalp event-related potentials (ERPs) exhibited a decreased amplitude over posterior electrodes for Repeat stimuli. Significant effects were observed during the late ERP component from 313 to 345 ms post-stimulus onset (all *p*s *< .*0028, *q* < .05; Fig. 1E). Similar ERP effects over posterior electrodes have been reported in prior EEG studies, although typically with much shorter delays between stimulus repetitions (e.g. ^25–28^).

**Fig. 1.**
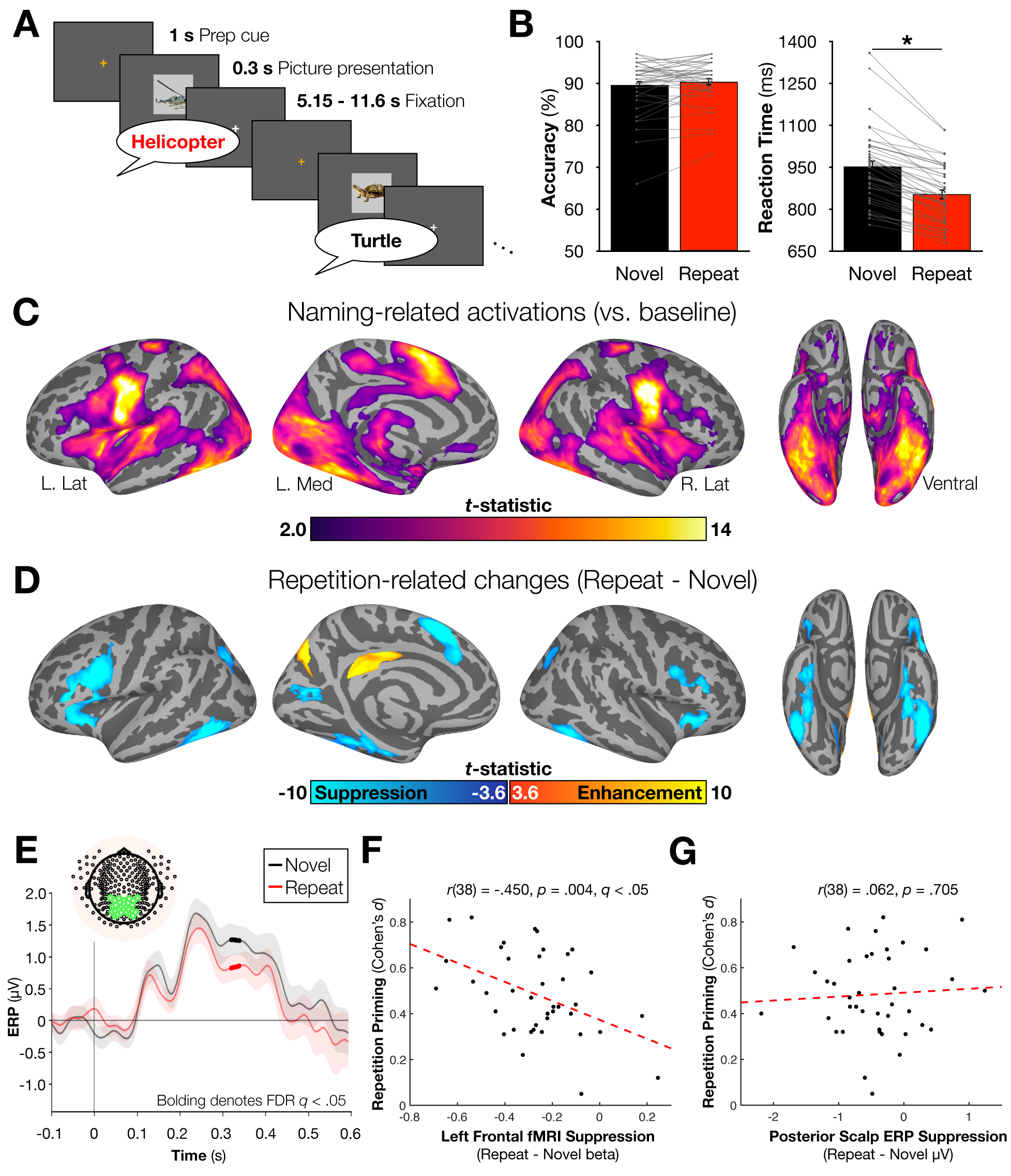
Behavioral, fMRI, and scalp ERP results associated with the object naming task (N = 40 participants). (A) The task utilized a slow event-related design. A preparatory (prep) cue was provided 1 s before each stimulus appeared. Stimuli appeared for 300 ms and were followed by a variable period of 5.15 – 11.6 s. Participants were instructed to speak the name of each image as quickly as possible. Novel and Repeat stimuli (items previously named approximately 75 minutes prior to brain imaging) were pseudorandomly intermixed. (B) Naming accuracy did not differ between the Novel and Repeat conditions, but participants demonstrated repetition priming by naming Repeat stimuli significantly faster than Novel stimuli. (C) fMRI identified significant activations in multiple regions across the cortex. (D) Within Naming-activated regions, significant repetition suppression was noted in multiple regions including bilateral frontal and fusiform cortex. Repetition-related increases in activity (repetition enhancement) were observed in the precuneus and mid-cingulate/rostral posterior cingulate cortex. (E) Significant repetition suppression of scalp ERP signals were observed over an *a priori*-defined posterior electrode montage. (F) Repetition priming was significantly correlated with fMRI repetition suppression in left frontal cortex, but (G) not with ERP suppression over the posterior electrode montage. * denotes p < .001.

To establish the relation between priming and RS in the current study, we first converted the priming scores for each participant to Cohen’s *d* effect sizes (following ^14^) and then correlated priming magnitude with RS in fMRI and in the scalp ERPs. Across participants, priming magnitude was significantly correlated with the magnitude of RS in left frontal cortex in fMRI, as has been observed previously^23^ [*r*(38) = -.450, *p = .*0036, *q* < .05; Fig. 1F], with non-significant correlations observed for the dACC/pre-SMA [*r*(38) = -.294, *p = .*065] and the left fusiform gyrus [*r*(38) = -.282, *p* = .078]. In contrast, RS in the scalp ERPs was not correlated with priming magnitude [*r*(38) = .062, *p* = .705; Fig. 1G], nor was it related to fMRI RS in any region (all *r*s < | .175 |, *p*s > .28). Partialling out scalp ERP RS also had little impact on the relationship between left frontal RS and priming magnitude [partial *r*(37) = -.447, *p* = .0043]. Taken together, these results suggest that left frontal RS in fMRI and RS in scalp ERPs index different aspects of neural activity, and it is the left frontal RS in fMRI that is most strongly associated with repetition priming.

### Oscillatory power rises and falls earlier with stimulus repetition in fMRI-localized regions of interest in a manner correlated with behavioral priming magnitudes

Whereas the analysis of scalp ERP RS enabled comparisons with prior relevant literature, the simultaneous acquisition of fMRI data allowed for a more targeted investigation of electrophysiological effects by examining source-estimated fluctuations of the EEG signal within fMRI-defined regions of interest. Trial-level EEG signals for dipoles perpendicular to the cortical surface were estimated from the scalp data for the four main activated regions showing RS in fMRI (left frontal, dACC/pre-SMA, left and right fusiform gyri; Fig. 2A) and decomposed into wavelets for time-frequency analyses (see *Methods*). Strong effects of stimulus repetition were observed in induced rather than evoked power (Supplementary Figs. 1-2) across the gamma (30- 80 Hz), beta (13-29 Hz), alpha (8-12 Hz) and theta (4-7 Hz) frequency bands in all four fMRI RS regions [all *t*s(39) > 2.206, *p*s *< .*0334, *q*s < .05; Fig. 2B]. These followed a stereotypical pattern, with an early increase in induced power for the Repeat relative to the Novel condition around 300-500 ms post-stimulus (earliest onsets detected in the gamma frequency range at 280 ms, 431 ms, 519 ms, and 385 ms in the left frontal, left fusiform, dACC/pre-SMA and right fusiform regions, respectively) through approximately 800-900 ms post-stimulus (latest offsets in the alpha frequency range at 888 ms, 870 ms, 892 ms, and 917 ms in the left frontal, left fusiform, dACC/pre-SMA and right fusiform regions, respectively; see bolded segments backed by light red background in Fig. 2C). This was followed by a more rapid decrease in induced power for the Repeat condition relative to the Novel condition around 1100-1200 ms post-stimulus (earliest onsets detected in the beta frequency range at 1171 ms, 1150 ms, 1081 ms, and 1112 ms in the left frontal, left fusiform, dACC/pre-SMA and right fusiform regions, respectively) through approximately 2100-2300 ms post-stimulus (latest offsets in the theta frequency range at 1888 ms, 2381 ms, 2138 ms, and 2609 ms in the left frontal, left fusiform, dACC/pre-SMA and right fusiform regions, respectively; see bolded segments backed by light blue background in Fig. 2C). However, no significant differences in the timing of effects were identified after correction for multiple comparisons. Additional effects in the delta frequency range (1-3 Hz) were also observed in the left fusiform [early: all *t*s(39) > 3.24, *p*s < .0025, *q*s < .05], dACC/pre-SMA [late: all *t*s(39) < -2.86, *p*s < .0068, *q*s < .05], and the right fusiform regions [late: all *t*s(39) < -3.05, *p*s < .0042, *q*s < .05] (Supplementary Figs. 1-2; periods of significance for each frequency and region are reported in Supplemental Table 1).

**Fig. 2.**
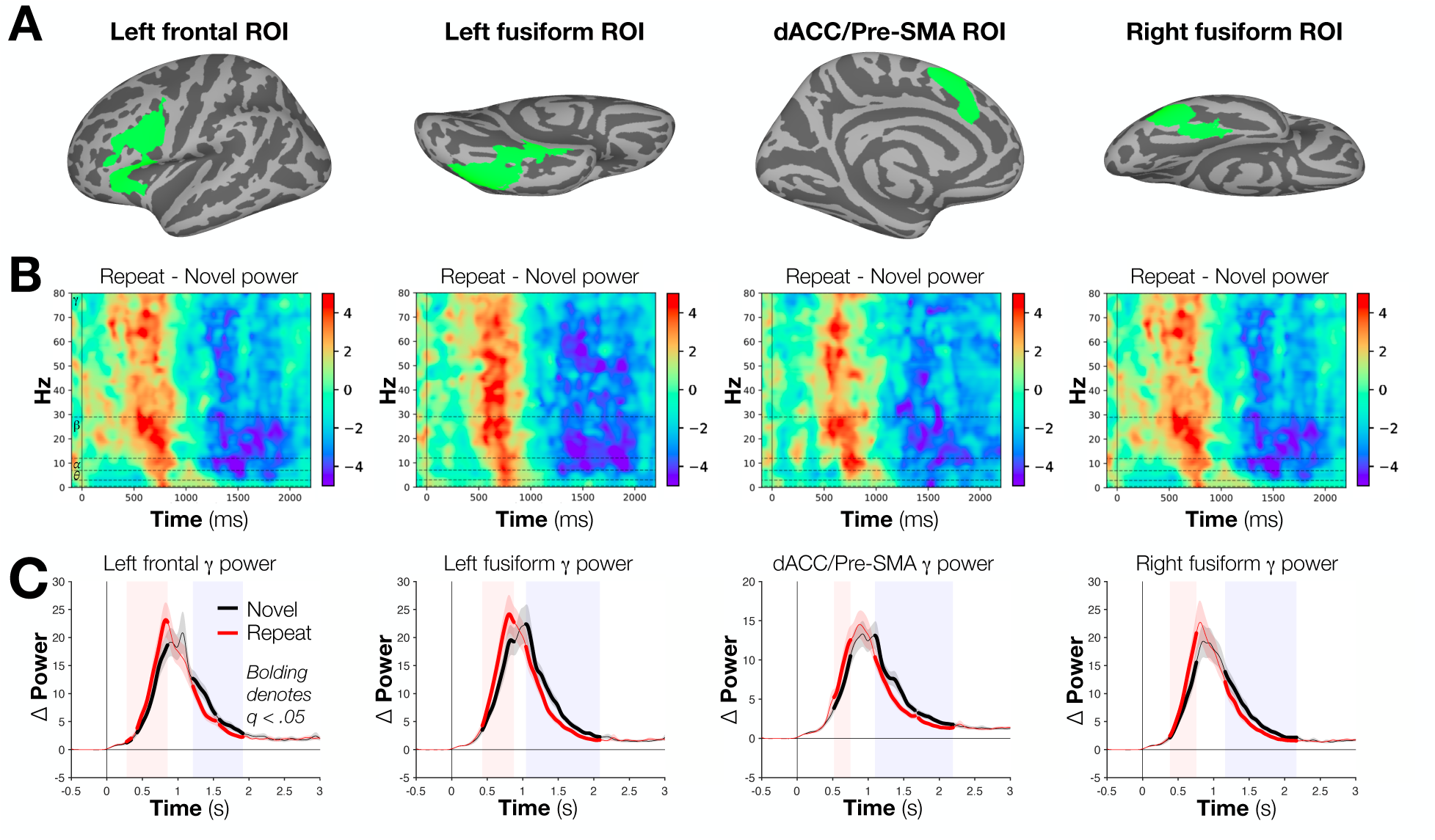
fMRI-identified regions of interest and induced power differences over time. (A) Regions of interest were identified in locations where masks of Naming-related activation (Fig. 1B) and Repetition Suppression (Fig. 1C) intersected at the group level (N = 40). (B) After source estimation of the EEG data, differences in Novel and Repeat conditions were plotted as a function of time and frequency in each region of interest. In all cases, a significant “early” period of Repeat > Novel activity was observed, with a “late” period exhibiting significant Novel > Repeat effects. (C) “Early” and “late” period differences can be appreciated by examining average activity in the gamma band. Bolded sections indicate periods of significant differences in power (FDR corrected to *q* < .05).

Having identified “early” repetition-related increases and “late” repetition-related decreases in induced EEG power, we then asked if the mean power differences during these periods correlated, across subjects, with either the magnitude of repetition priming or of BOLD RS in the four fMRI-defined regions of interest. A significant (Spearman) correlation was identified between differences in local EEG power and fMRI RS in the left frontal region for the early period (Repeat > Novel) in the gamma band [*r*(38) = -0.37, *p < .*0189, *q* < .05; Fig. 3A]. In contrast, correlations between differences in EEG power and priming magnitude across participants were observed broadly across the four regions in both the early and late periods for the gamma and beta bands [*r*s(38) > | 0.3906 |, *p*s *< .*0128, *q*s < .05 for all] and in the late periods for the alpha and theta bands, as well [*r*s(38) > | 0.4197|, *p*s *< .*008, *q*s < .05 for all] (see Fig.3B- E for left frontal correlations; see Supplementary Table 2 for all results by region and frequency band).

**Fig. 3.**
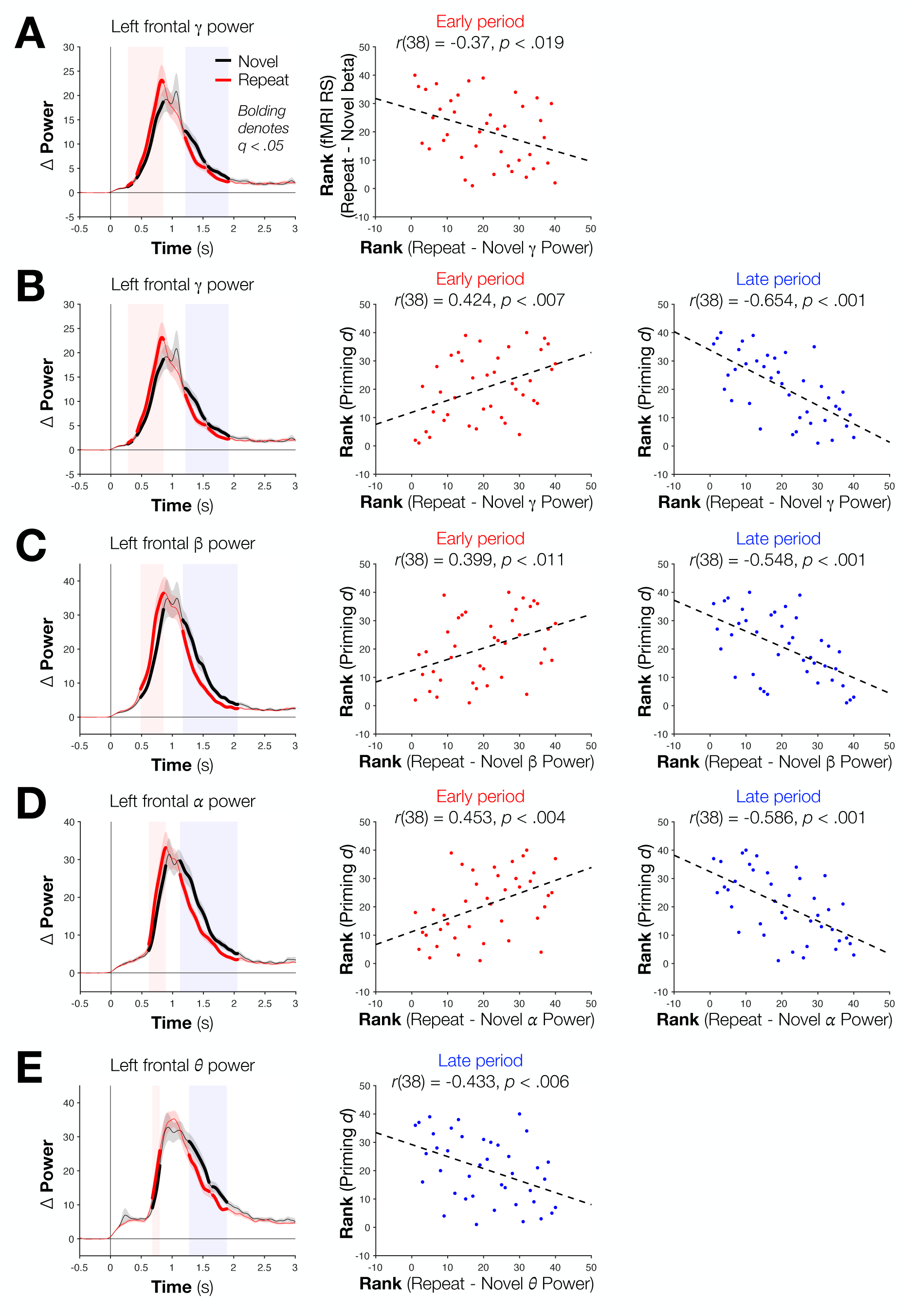
Left frontal gamma and beta power differences correlate across participants (N = 40) with fMRI repetition suppression in the same region and with behavioral priming. (A) Significant Repeat > Novel gamma power was observed in an early period, whereas significant Novel > Repeat power was observed in a late period. Across participants, EEG power differences during the late window and fMRI repetition suppression magnitudes were significantly (Spearman) correlated. (B) Within the gamma band, early periods of Repeat > Novel power differences, and later Novel > Repeat differences, both correlate (Spearman) with the magnitude of repetition priming. (C) This same pattern was observed in the beta band, as well as (D) in the alpha band. (D) “Late” window correlations were also observed in the theta band.

The pattern of power increases and decreases for Repeat versus Novel conditions shown in Figures 2 and 3 suggested a leftward shift of the curves for the Repeat condition to earlier time points, rather than a more complex difference in overall magnitude of the curves. Importantly, such a temporal shift is predicted by the Facilitation model. To test this issue, we examined the sum of the area under the power curves for the two conditions across participants. The curves did not significantly differ in any band or region following correction for multiple comparisons, although three uncorrected differences (Sign Test, all uncorrected *ps < .*039 for Novel > Repeat) were observed for beta power in the frontal and right fusiform regions and for theta power in the dACC/pre-SMA region. These results indicate that the large “early” and “late” differences between conditions predominantly reflect the shifting of the Repeat curves to earlier timepoints and not differences in the overall amount of power relative to a pre-stimulus baseline. This shift can also help to explain the change in signs between early and late effects—the former is due to the more rapid rise in power for Repeat items, and the latter is consistent with a more rapid fall for those same Repeat items.

In the left frontal region, the “early” power difference in the gamma frequency was jointly correlated with both fMRI RS and priming. We therefore examined whether these effects share a common underlying pool of variance or reflect unique variances using partial correlation. The correlation between power differences and priming magnitude remained after partialling out left frontal fMRI RS magnitude [partial *r*(37) = .335, *p* = .037]. In contrast, the correlation between left frontal fMRI RS and priming magnitude—a well-replicated finding in this literature^11,13,23^—was no longer significant after partialling out power differences [partial *r*(37) = -.24, *p* = .14]. EEG induced power differences can therefore explain unique variance in priming magnitude beyond that explained by fMRI RS, but fMRI RS, in turn, fails to explain significant variance beyond what is shared with induced power differences.

### Oscillatory power differences and their correlations with behavior do not reflect the overt speech response

It was possible that the differences in power between Repeat and Novel conditions or their correlations with behavior could have been due to speech-related artifacts, such as an electromyographic (EMG) response in articulatory muscles. This possibility was investigated in several ways. First, results were compared with and without the ICA-based EMG cleaning of the data. Although the amplitudes of the ERP waveforms were decreased by the ICA-based removal of the components during the time of the overt responses that explained the top 90% of time series variance, the correlations between the induced power differences and behavior marginally increased in magnitude when the cleaning was applied (*p* < .09, permutation test; Supplementary Figs 3-4, Supplementary Table 3). Second, we examined whether correlations between power differences and priming magnitudes were strongly altered by controlling for mean response times (which are, necessarily, coupled to the speech/motor response) through partial correlation. All of the initially detected correlations remain significant when response time was controlled in this manner (see Supplementary Table 4). For example, the across-subjects correlation (Spearman) of the “early” power differences in the left frontal region with priming was *r*(38) = 0.4236, *p = .*0065 before, and *r*(37) = 0.4239, *p = .*0072 after, partialling out mean response time across participants. Third, we checked whether our source-estimated power results at each node on the cortical surface were consistent with an EMG or motor response origin. Again, these analyses failed to indicate an artifactual basis to the observed results (see Supplementary Movies 1-4). We also back-projected the estimated cortical sources to the scalp to identify the most-important scalp electrodes for each source region (Supplementary Fig. 5). This provided further evidence that the four cortical surface regions were not due to a common spatial artifact and permitted companion analyses on spatially distinct collections of scalp electrodes (Supplementary Figs. 6- 9).

Finally, we prospectively designed and conducted an additional experiment (*N* = 25) in which participants covertly named pictures during simultaneous fMRI-EEG, indicating when they had identified each object via a button press in their left (non-dominant) hand (see *Methods*). We used the time windows of significance and fMRI regions of interest from the overt naming experiment to evaluate any power differences and correlations with priming magnitude. As expected from prior studies^23,29^, priming was somewhat attenuated during covert naming [Repeat *M*(SD) = 572.2 (74.9) ms, Novel *M*(SD) = 642.1 (107.5) ms; *t*(24) = 6.47, *p < .*001, *d* = 1.29]. However, fMRI RS was still observed in the same four regions of interest identified in overt naming [left frontal *t*(24) = 5.97, *p* < .001, left fusiform gyrus *t*(24) = 3.78, *p* < .001, dACC/pre-SMA *t*(24) = 8.12, *p* < .001, right fusiform gyrus *t*(24) = 2.78, *p* = .01; Fig. 4A, see also Supplementary Fig. 10]. A significant correlation between fMRI RS and priming magnitude was observed in the left fusiform region [*r*(23) = -.405, *p* = .044, and a similar but non-significant correlation between fMRI RS and priming was observed in the left frontal region (*r* = -.386, *p* = .057; Fig. 4B). In all four regions, time-frequency analysis identified “early” periods of greater power for Repeat than Novel stimuli in at least one frequency band (left frontal theta power: all *t*s > 2.64, *p*s < .0143, *q*s <.05; left fusiform gamma power: all *t*s > 2.55, *p*s < .0175, *q*s < .05; left fusiform alpha power: all *t*s > 2.77, *p*s < .0106, *q*s < .05; dACC/pre-SMA theta power: all *t*s > 2.96, *p*s < .0069, *q*s < .05; right fusiform theta power: all *t*s > 2.74, *p*s < .0115, *q*s < .05; Fig. 4C,D). One “late” period of greater power for Novel than Repeat stimuli was also detected for the right fusiform region in the theta frequency range (all *t*s < -2.74, *p*s < .0115, *q*s < .05;Fig. 4D, rightmost column). While effects were not equally detected in all frequencies, the overall qualitative pattern of time-frequency power differences appears consistent with those seen in overt naming, with relatively broadband early and late bands at approximately the same time periods.

**Fig. 4.**
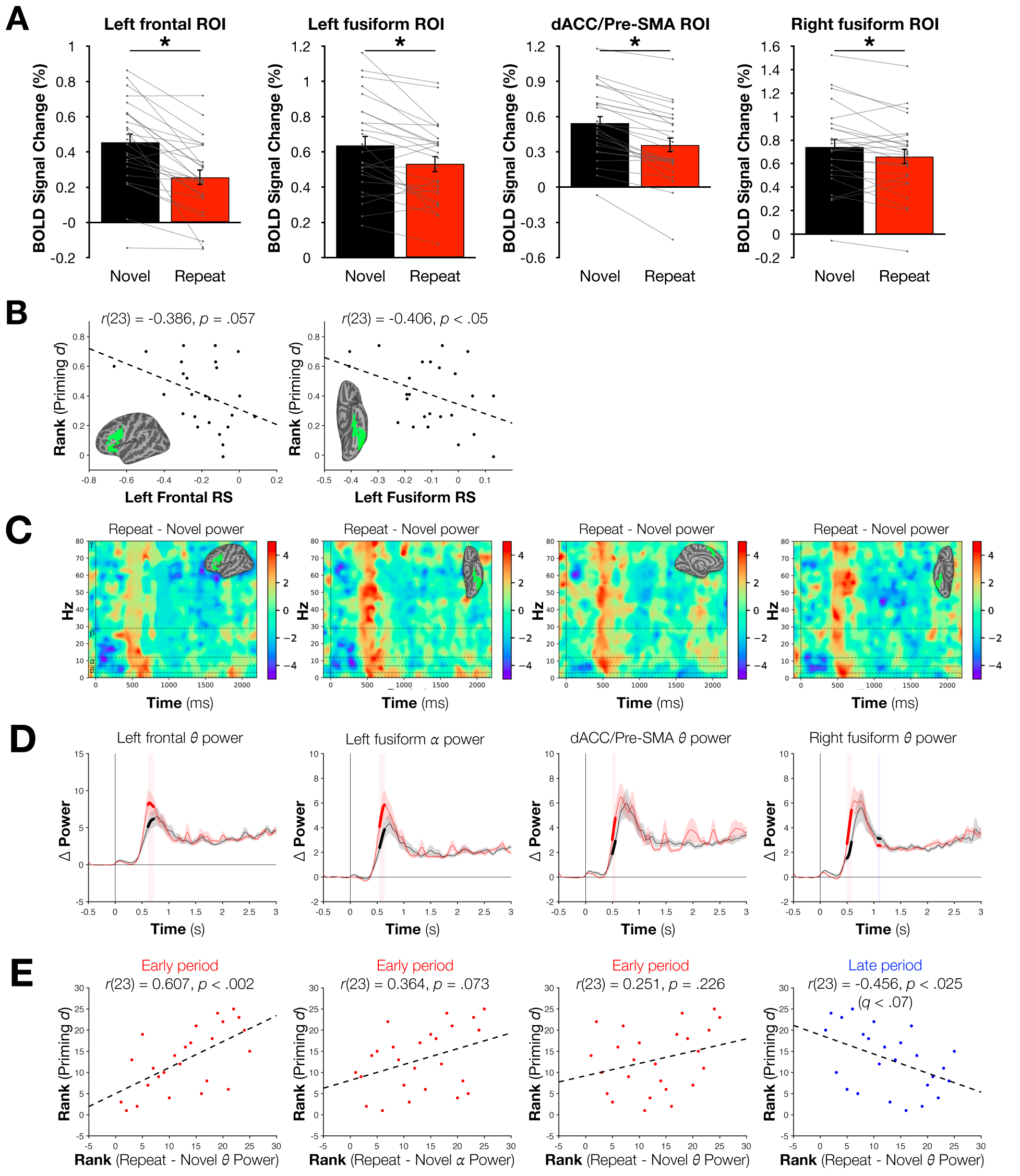
Results associating repetition priming with fMRI repetition suppression and differences in EEG power replicate when no spoken responses are required. (A) Within each of the 4 regions of interest defined in the overt naming data, significant repetition suppression effects were observed in a “covert naming” experiment (N = 25). (B) The magnitude of repetition suppression effects was significantly correlated (Spearman) with repetition priming across subjects in the left fusiform region, with a similar but nonsignificant correlation observed in left frontal cortex. (C) An early period of Repeat > Novel power was observed across in time- frequency analysis of EEG data in every region, (D) most commonly in the theta or alpha bands. (E) The magnitude of “early” Repeat > Novel theta power correlated (Spearman) significantly across participants with repetition priming in the left frontal cortex, as did “late” theta power differences in right fusiform cortex (albeit at an uncorrected level). * denotes *p* ≤ .01.

As in the overt naming experiment, we tested whether the corrected differences in induced power between the Repeat and Novel conditions were correlated with fMRI RS and priming. The early difference in theta frequencies in the left frontal region was significantly correlated (Spearman) with priming magnitude [*r*(23) = .607, *p* < .0014, *q* < .05] , and the late difference in theta frequencies in the right fusiform region was correlated with priming magnitude at an uncorrected level [*r*(23) = -.456, *p* < .0219, *q* < .07] (Fig. 4E). No significant associations with fMRI RS were detected. Taken together, these findings demonstrate that the early and late power differences seen in overt naming are also observed in covert naming and therefore cannot be due to overt speech artifacts. Furthermore, priming is also associated with induced power differences in covert naming, although with the largest effects seen in somewhat lower frequencies (theta) than in overt naming (most robust in gamma/beta). Future work can clarify the nature of this observed difference.

## Discussion

In summary, we have found that repetition-related improvements in behavior (priming) and decreases in neural activity (repetition suppression) during object identification are accompanied by more rapid increases (and subsequent decreases) in oscillatory power in the repeated condition in concurrently acquired EEG. Power changes were robustly correlated with the magnitude of priming across participants, overlapping in variance explained with fMRI RS, while also explaining unique additional variance. Results, particularly in left frontal cortex, were qualitatively replicated when using a covert naming task, ruling out an artifactual basis of the power and priming relationships and lending support to their robustness.

Our results suggest that priming may be related to changes in neural activity through a novel combination of the Synchrony and Facilitation models. The Synchrony model predicts enhanced population-level oscillatory power when processing repeated items, as was observed here in the “early” period. Facilitation predicts more rapid onsets and offsets, although the current data diverge from earlier articulations of Facilitation^18,19^ insofar as the differences were observed in the domain of oscillatory power rather than total neural activity *per se*. However, facilitation also provides a means of accounting for both early and late periods of repetition-related differences in activity: given an earlier rise in and resolution of processing Repeat items, one would expect an early period of Repeat > Novel power differences due to the leftward shift of the power curve (Fig. 2). This same shift would then produce a later period in which an opposite pattern would necessarily emerge as the processing of Novel items is slower to resolve. It is therefore the early period that seems most relevant for understanding the relation between repetition priming and repetition suppression. Notably, although we have documented these changes across all four of the fMRI-defined regions exhibiting RS, prior work has highlighted a particularly critical role of left frontal cortex in mediating both repetition priming and repetition suppression across changes in task and stimulus form, implicating response learning mechanisms^13,15,30^. We do not perceive any incompatibility between these proposals and our current results. Indeed, one view is that this novel combination of Synchrony and Facilitation models is reflecting stimulus-to-response mapping processes.

Consistent with the present findings, prior EEG studies with repeated pictures have shown differences in induced gamma power^26,28^, although these have often utilized much shorter-term repetitions, sometimes without a requirement of overt responses to the stimuli of interest^26^. One prior study^31^ notably identified increased induced gamma power for well-known objects relative to novel objects in addition to the EEG RS effects that we also show, replicating this effect across two settings in two different labs. Similar results for increased gamma power and coherence with stimulus repetition have also been observed with microelectrode recordings in monkey V1 and V4 simultaneously with overall decreases in firing rate to grating and natural object stimuli^32–35^. Repetition-induced decreases in firing rate that coincide with increases in synchronous activity have even been observed in insects such as locusts^36^ and drosophila^37^, suggesting a relatively general mechanism of neural processing efficiency that has been preserved across species.

The pattern of results seen in overt picture naming was replicated qualitatively in covert naming for “early” (Repeat > Novel power in all four regions showing fMRI RS) and “late” periods (Novel > Repeat power in the right fusiform region). We further detected a correlation between early power differences in the left frontal region and priming magnitude, with an uncorrected correlation for the late period in the right fusiform region. While the early/late pattern appeared to be relatively broadband, the correlations with priming were observed in theta rather than gamma frequencies. These differences may be due at least in part to the smaller effect sizes typically seen in covert naming^23^, as well as the smaller sample size utilized in the covert experiment (covert *N* = 25; overt *N* = 40). The covert naming experiment was designed primarily to evaluate the continued presence of the early/late power difference pattern in the absence of overt speech rather than to replicate the correlations with behavior, and future studies with this and other tasks should be examined with larger samples to assess any real task-specific differences in the neural correlates of priming.

By linking the activity states predicted by the Synchrony and Facilitation models to the magnitude of both RS and priming, our findings provide a mechanistic explanation for the long-standing puzzle of how reduced neural activity can support behavioral improvements rather than behavioral impairments—a relationship that has been observed across humans, monkeys, and rodents. While the population-level measures of neural activity that are utilized in non-invasive human neuroimaging cannot yield direct evidence of the synchronization of neural firing, the family of effects shown using the current multimodal approach is consistent with earlier, enhanced neural synchronization, and it provides an important target for future work using more direct neural recording techniques in animals and human patients.

## METHODS

### Participants

#### Overt Naming

A total of 45 participants were recruited from the National Institutes of Health and surrounding area. Five of these participants were excluded (1 following detection of a neurological abnormality, 1 for excessive motion in 75% of task scans, 2 due to equipment malfunctions, and 1 because their vision could not be corrected with MR compatible lenses). The remaining 40 participants (18 male) had a mean age of 24.4 ±2.9 years (range: 19-32).

Participants had a mean of 16.2 ±0.97 years of education (range: 14-19), were right-handed, had normal or corrected-to-normal vision, reported no history of psychiatric or neurologic illness, and were native speakers of English. Informed consent was obtained from all participants and the experiment was approved by the NIH Institutional Review Board (Protocol 93-M-0170; clinical trials number NCT00001360). Participants received monetary compensation for their participation.

#### Covert Naming

Twenty-six participants were recruited from the National Institutes of Health and surrounding area. One participant was excluded when sufficiently good contact could not be made between EEG sensors and their scalp. The remaining 25 participants (10 male) had a mean age of 26.4±4.3 years (range: 21-35) and had completed 16.6±1.15 years of education (range: 14- 19). Participants were right-handed, had normal or corrected-to-normal vision, reported no history of psychiatric or neurologic illness, and were native speakers of English. Informed consent was obtained from all participants and the experiment was approved by the NIH Institutional Review Board (Protocol 93-M-0170; clinical trials number NCT00001360). Participants received monetary compensation for their participation.

### Experimental stimuli

Stimuli consisted of 200 colored photographic images of animals, plants, and everyday objects, and have been used in prior laboratory studies (e.g., ^23,24^). Images were split into 2 groups (lists) of 100 images each, which were matched in average lexical properties including log HAL frequency (means = 8.54 vs. 8.60, *t*(198) = .293, *p* = .770), name length (5.55 vs. 5.93 letters, *t*(198) = 1.31, *p* = .191), and average naming time based on prior studies in the laboratory using a similar paradigm (824.8 ms vs. 831.8 ms, *t*(198) = .496, *p* = .621). Experimental task scripts were written in PsychoPy2 software. In-scanner stimulus presentation was accomplished using a desktop running Windows 10 connected to a 120 Hz, 1920x1080 pixel MR-compatible monitor. Images subtended approximately 5° of visual angle and were centrally presented against a dark gray background. Participants viewed the monitor through a mirror attached to the head coil. A central white fixation cross was shown between stimulus presentations. Out-of-scanner stimulus presentation was conducted using a Macbook Pro laptop (Apple, Cupertino, CA) in a behavioral testing room.

### Task Design

#### Pre-scan object naming

Prior to scanning and EEG cap placement, participants in both Overt and Covert Naming tasks viewed and overtly named 100 images twice through in a testing room. Images were from one of the predetermined lists described previously, and the list used during this portion of the task was counterbalanced across participants. All images were presented a single time before any were re-presented. 500 ms prior to each image presentation, the white fixation cross changed to an orange color to serve as an onset preparation cue. Images were then presented for 300 ms and followed by a white fixation cross for 1700 ms. Participants were instructed to name aloud each image as quickly, accurately, and clearly as possible, and their responses were logged by an experimenter. A schematic of this task structure is presented in Fig. 1A.

Following the pre-scan object naming phase, participants were fitted with an MRI-compatible EEG cap. Electrode locations and impedances were then recorded and adjusted as necessary (described in more detail under “EEG data acquisition”), and participants were then led into the scanner to complete the in-scanner object naming task. The average delay between the end of the pre-scan object naming period and the beginning of in-scanner object naming was approximately 75 minutes.

### In-scanner object naming

Participants underwent simultaneous EEG-fMRI scanning while again naming objects.

200 items were now presented in total: 100 were repeated from the pre-scan naming phase (Repeat), and 100 newly presented images (Novel). Repeat and Novel images were intermixed throughout each scan run. The basic trial structure remained similar but was modified to fit a slow event-related fMRI design with widely spaced trials (Fig. 1B). 1000 ms before each image appeared, the fixation cross changed to an orange color to serve as an onset preparation cue.

Images were then presented for 300 ms and followed by a white fixation cross for a variable period of 5150-11600 ms (in steps of the TR = 2150 ms). The order of stimuli and the delay period following each stimulus were pseudorandomized to maximize design efficiency^38^. Overt Naming participants were instructed to name aloud each image as quickly, accurately, and clearly as possible. Responses were recorded for subsequent scoring and response time (RT) calculation (see “Behavioral data acquisition” below). Covert Naming participants did not speak words aloud but instead pressed a button with their non-dominant hand (left) to indicate the beginning of their naming response (as in ^23,29^).

### Post-scan object naming (Covert Naming participants only)

Participants completing the Covert Naming task in the scanner also completed an additional post-scan session in which they named all 200 pictures immediately after completion of fMRI scanning and the removal of the EEG net. This allowed for overt naming of all types of stimuli, and any stimuli that were not successfully named during the post-scan session had their associated in-scanner trials marked as incorrect.

### Data acquisition

#### fMRI data acquisition

Scanning was conducted on a General Electric Discovery MR750 3.0T scanner, using a 32-channel head coil. Separate high-resolution T1 structural images were obtained for each participant both with the EEG cap and without it (TE = 3.47 ms, TR = 2.53s, TI = 900 ms, flip angle = 7°, 172 slices with 1×1×1mm voxels). For 19 subjects, T1 scans without the cap were not collected on the day of scanning but were instead obtained in separate sessions. Functional images were acquired using a BOLD-contrast sensitive multi-echo echo-planar sequence (Array Spatial Sensitivity Encoding Technique or ASSET acceleration factor = 2; TEs = 13.5, 34.2, 54.8, and 75.5 ms; TR = 2150 ms, flip angle = 75°; 72 × 72 matrix, in-plane resolution = 3.0 mm).

Whole brain EPI volumes (MR frames) consisted of 42 interleaved, 3.0 mm-thick oblique axial slices, aligned manually to the AC-PC axis. One additional EPI scan, with identical parameters but an inverted phase encoding direction (P-A), was collected prior to task scans for purposes of distortion correction during preprocessing. Foam pillows helped stabilize head position for all participants, and foam earplugs attenuated scanner noise. A sensor was placed on each participant’s left middle finger to record heart rate and a respiration belt monitored breathing for each participant.

#### EEG data acquisition

EEG data were collected using a 256-electrode (Ag/AgCl) HydroCel GSN 220MR Geodesic Sensor Net (Magstim EGI, Plymouth, MN, USA). Three different sized nets were available to accommodate different head sizes, but all were structured identically. Participant head circumferences were measured to identify the appropriate net size. Prior to application, each net was soaked in warm water containing a combination of potassium chloride and baby shampoo. Placement indicators were marked on each participant’s head with a watercolor pencil to signify the Cz point. Following placement of the net, the specific positions of each electrode were recorded using a handheld EGI GeoScan stereo camera and its accompanying software.

Impedances were measured using EGI NetStation v5.4.2 and experimenters made adjustments until electrical impendences were below 40 kΩ. Once completed, a tubular elastic net was placed over the EEG net to stabilize the electrodes prior to participants entering the scanner.

EEG signals were digitized at a 1000 Hz sampling rate with a Net Amps 400 amplifier and recorded onto a MacBook Pro laptop (Apple Inc., Cupertino, CA) running EGI NetStation. Data were collected unfiltered. Horizontal eye movements (i.e., saccades) and blinks were monitored using built-in electrodes on the sensor net^39^. EEG data were recorded using a Cz reference. For 20 overt naming participants, two additional sensors were placed on the chest for collecting EKG data (hardware to allow this for the first 20 participants was not available due to COVID-related supply chain disruptions). EKG was recorded with the same parameters as the EEG signal (unfiltered, digitized at 1000Hz).

The EEG system clock was synchronized with and slaved to the MRI scanner clock (10 MHz polling rate), and synchronization was further improved by collection of sync pulses that were sent from the scanner at the start of each TR. Orange fixation cross and object picture onset times were sent from the stimulus PC to the NetStation recording computer using PsychoPy2’s *egi* package (https://psychopy.org/api/hardware/egi.html). A 37 ms asynchrony (average delay) was measured with a photodiode between picture onsets on the monitor and the signal sent to NetStation; this was subsequently accounted for in EEG data analyses.

#### Behavioral data acquisition

Participants spoke all responses into a FOMRI-III NC MR-compatible microphone (OptoAcoustics Ltd., Mazor, Israel). Audio signals from this microphone were routed into an M-Audio FastTrack Ultra 8-R USB audio interface, which in turn was connected to a Dell Precision M4400 laptop. Responses were recorded using Adobe Audition. In conjunction with the spoken audio recording, the stimulus presentation computer sent out a square wave pulse at the onset of each stimulus that was captured on a parallel audio track by the recording laptop. Onset time differences between the square pulse onset and the voice response in each trial were used to calculate voice RTs^23,24^. Naming responses were recorded by the experimenter and subsequently marked as correct/incorrect.

### Data preprocessing

#### MRI data preprocessing

MRI data were preprocessed using AFNI^40^, Freesurfer^41^ v7.2, and SUMA^42^. Dicoms were converted to 3- (for structural scans) or 4-dimensional (for functional scans) files using *dcm2niix_afni*. T1 structural images were then processed using Freesurfer’s *recon-all* tool, skull-stripped using *@SSwarper*, and standard SUMA std.141 surface meshes for each participant were generated using *@SUMA_Make_Spec_FS*. The first 4 MR frames of each EPI scan run were excluded to allow for T1 equilibration. Slice-time correction was subsequently carried out for each echo of each run. Blip-up/blip-down distortion correction was carried out using AFNI’s *unWarpEPI.py* script for each run using the scan run with an inverted phase encoding direction. EPI data were realigned to the first volume of each scan run via rigid body transformation and subsequently registered to the anatomical image. Motion was calculated using the second echo of each run and motion estimates were retained for use as nuisance regressors during subsequent general linear model (GLM) analysis. Functional data then underwent multi-echo ICA analysis using *tedana.py* software^43,44^. This procedure initially calculates a weighted average of the different echo times within each voxel and then uses spatial ICA and the known properties of T2* signal decay over time (i.e., echoes) to separate and reject putatively artefactual components, such as thermal noise or head motion, while retaining components consistent with the Blood Oxygen Level Dependent (BOLD) response. Denoised timeseries data were mapped to each participant’s std.141 surface mesh and a geodesic blur along the unfolded cortical surface was applied to achieve a FWHM of 8 mm. Data were then normalized and rescaled to percent signal change.

All task scan runs consisted of 237 MR frames (233 after initial frame discarding) and were approximately 8 min, 30 s in length. Four naming scans were completed for each participant. Average run-level motion estimates were derived using AFNI’s *@1dDiffMag* based on three translational and three rotational motion parameters; runs with values > 0.4 mm/TR were excluded. For the final Overt Naming sample, 2 participants had 2 runs excluded based on this requirement. For the Covert Naming sample, a single participant had 2 runs excluded for excessive motion. For all retained runs, single TRs were censored if frame-to-frame motion exceeded a Euclidian Norm of 0.3 mm or if ≥ 10% of voxels in that TR were flagged as outliers by AFNI’s *3dToutcount*. This resulted in an average of 6±2% of TRs being censored for Overt Naming participants and 5±1% for the Covert Naming participants.

### EEG data preprocessing

EEG data were initially processed using NetStation. First, MR-induced gradient artifacts were removed from the EEG using an average artifact subtraction based on a sliding window of 10 TR pulse events. Ballistocardiogram (BCG) artifacts were then removed. For the participants for whom EKG sensors were not available, BCG effects were estimated from the EEG data using NetStation’s built-in BCG identification algorithm; for subjects for whom EKG was collected, QRS complexes of the heartbeat were defined in the EKG channel. In both cases, BCG removal was accomplished using an optimal basis sets approach^45^.

Two pipelines were used to pre-process EEG data following removal of MR gradient and BCG artifacts (Supplementary Fig. 4A). Both interpolate bad channels and attenuate electrooculographic (EOG) and tongue-related electromyographic (EMG) artefacts. The second pipeline includes an extra step to attenuate speech-related EMG. Both pipelines were fully conducted using MNE-Python version 1.8 (stable)^46^.

### Band-pass filtering and handling of bad channels

EEG data were re-referenced to the Cz channel (referred to as “VREF” by the EGI software) and bandpass-filtered from 0.1 to 100 Hz using *mne.Raw.filter* with default arguments. Next, the 60 Hz frequency band and its integer multiples were notched out using “mne.filter.notch filter” with default arguments, except for the phase argument that was set to “zero-double”. The filtered data were then epoched (broken down into trials) ranging from -3 to 5 seconds relative to stimulus onset time. Finally, each individual trial was screened for “bad channels”, that is, channels with peak-to-peak amplitude > median + 1.5 interquartile range. Bad channels were interpolated using *mne.Epochs.interpolate bads* with default arguments.

### Attenuation of EOG and tongue EMG

EEG activity that resulted from eye, eyelid and tongue movements was removed from each channel of each trial by regressing out activity measured at electrodes at the left and right superior orbicularis oculi (in the EGI-256 montage, channels E37 and E18, respectively), left and right inferior orbicularis oculi (E241 and E238) and those at the outer cantus left and right (E252 and E226)^39^. Activity in these channels was regressed out using *mne.preprocessing.EOGRegression* with the argument “picks_artifact” set to these electrodes.

The outputted weights were then used (*weights.apply*) to generate the denoised EEG. To avoid regression overfitting, the coefficients of determination (R^2^’s) of all regressions/channels were inspected. Channels with an R^2^ < 0 were treated as bad channels and interpolated using *mne.Epochs.interpolate bads* with default arguments.

### Attenuation of speech EMG

An additional denoising step was also applied to each channel for each trial. The purpose was to mitigate putative EMG artefacts due to contraction of articulatory muscles during speech preparation and production. First, for a given trial, a time window of -0.2 to 0.2 s was centered at the RT, and the corresponding electrode data were fed to Independent Component Analysis (ICA) yielding speech independent components (ICs). This was accomplished using *mne.preprocessing.ICA*, specifying “method”=picard and “n components”=0.9. Note that, in MNE Python, ICA is always preceded by Principal Component Analysis (PCA). By setting “n components”=0.9, the algorithm constrained ICA to operate on the minimum number of principal components that spanned 90% of the variance. Next, *mne.preprocessing.EOGRegression* was applied using the argument “picks_artifact” set to the time series of speech ICs. The outputted weights were then used (*weights.apply*”) to generate additionally denoised timeseries data. Note that while the ICA was conducted on the period around the RT (-.2 to +.2 sec), the weights were obtained from and applied to a window from 0.5 s prior to the RT up until the RT, a period of the trial considered representative of speech preparation and production. As with attenuation of EOG and tongue EMG, regression overfitting was avoided by treating channels with an R^2^ < 0 as bad channels and interpolated using *mne.Epochs.interpolate bads* with default arguments.

### Data analysis

#### Behavioral analysis

Repetition priming was defined as the mean of the naming response times (Overt Naming) or the mean of the button press times (Covert Naming) for the Novel condition minus the mean of the response times for the Repeat condition. Only correct trials were used for these calculations. Repetition priming magnitudes were then estimated for each participant using Cohen’s *d* (effect size) in order to adjust for different response time ranges for each participant (see ^14,23^ for discussion). Accuracy (percentage of correct responses) in the Novel and Repeat conditions was also calculated for each participant. Statistical analyses were then carried out across participants using paired *t*-tests for accuracy and repetition priming in both Overt Naming and Covert Naming.

#### fMRI data analysis

fMRI data were analyzed using a GLM approach. Responses for each condition were modeled using TENT functions, which is analogous to SPM’s finite impulse response (FIR) model. This approach assumes that all responses for a given condition share the same response shape but makes no assumption as to what the shape of that response might be. Ten time points, extending from 0 s to 19.35 s inclusive in steps of the TR (2.15 sec), modeled each of 3 conditions of interest. These were Repeat trials (for images named during the pre-scan period as well as fMRI scanning), Novel trials (images named only during the fMRI scanning period), and Error trials. Error trials included multiple types of errors, including non-responses, incorrect responses, responses for which responses changed between the pre-scan and in-scanner naming periods (Repeat trials only), or trials in which responses were provided after a 2150 ms (1 TR) delay (as in ^23^). For the purposes of statistical testing, response magnitudes were estimated for each condition by averaging the 3rd and 4th time points of the TENT function (reflecting the expected peak of the BOLD response, 4.3 to 8.6 s following stimulus onset).

Two vertexwise contrast maps were generated from the fMRI data. First, a contrast (paired-samples, two-tailed) of the regression coefficients (beta weights) for the Repeat – Novel conditions identified vertices for which significant repetition suppression (Repeat < Novel) or repetition enhancement (Repeat > Novel) effects were observed. Correction for vertex-wise multiple comparisons was accomplished using the *SurfClust* function in AFNI, which uses the smoothness of the data to generate null surface maps and empirically defines a minimum cluster size to achieve a given whole-brain correction at a given significance level. A vertex-wise threshold of *p* < .001 and a minimum cluster extent of 195 vertices achieved a corrected *p* < .05 in this “Repetition-related changes” map.

A second contrast (one sample, two-tailed) was conducted to identify vertices generally activated by the viewing and naming of picture stimuli, starting from the moment of image onset (irrespective of sensitivity to repetition-related effects). For each vertex, the beta weights for Repeat and Novel conditions were first averaged and then tested vs. a baseline of 0. Significant vertices (FDR corrected to *q* < .05) were included in this “Naming-related activation” map.

The Naming-related activation map was then used as an inclusive mask to identify cortical regions that were both activated by naming and that exhibited significant repetition-related changes in activity. Regions of interest (ROIs) were identified in left frontal cortex, left and right fusiform cortex, and the dACC/pre-SMA. Prior work using this paradigm and approach to fMRI data analysis^23^ identified clusters in these regions, and the largest clusters in the current masked data that corresponded to each location were manually identified and carried forward for further analysis. The magnitude of repetition suppression in these ROIs was then calculated by averaging the Repeat-Novel beta weights across all vertices associated with each region, respectively, for each participant.

### EEG scalp analysis

Following prior findings of repetition-related EEG effects over posterior locations^31^, our scalp analyses focused on posterior electrodes. In the EGI montage, these consisted of electrodes E108, E109, E115, E116, E117, E123, E124, E125, E138, E139, E140, E149, E150, E158, E159, E87, E99, E110, E118, E126, E100, E119, E127, E101, E128, E129, E141, and E153. Repetition suppression was defined as the ERP difference between Novel vs. Repeat conditions. Posterior ERPs were calculated as an average timeseries across trial epochs (-0.5 to 3.0 s relative to stimulus onset) at the selected electrodes, with a separate ERP calculated for each condition (Novel, Repeat) and participant. The mean of the baseline period (-.5 to 0 s relative to stimulus onset) was subtracted from each trial epoch prior to averaging. The mean voltage of the Novel – Repeat ERP waveforms was compared with paired *t*-tests across participants at each time sample, with multiple comparisons controlled by False Discovery Rate (FDR, *q* < .05). The time period used for FDR correction was 0.0 to 0.6 s to match the analysis window of prior related work^31^.

### EEG source analysis

#### Forward solution

For each participant, we used Freesurfer’s *recon-all* to derive 3-D surfaces from that participant’s T1 MRI images. Next, we used *mne.setup source space* with the spacing argument set to “oct6” to define a source mesh over the participant’s white matter, with sources spaced by 4.9 mm. To model current conduction, we set up boundary element models (BEMs) with 5120 dipoles per hemisphere. We first ran *mne.make bem model* with argument ico=4 and default conductivity arguments to generate the surfaces used by *mne.make bem solution*. This function was run on the brain, inner skull and outer skull surfaces output by Freesurfer’s *mri watershed* segmentation utility. The resulting BEM was used when generating each participant’s forward model. We obtained the forward model using *mne.make forward solution*. We set the head-fMRI transform argument “trans” to “fsaverage” to calculate the transformation between the EEG montage sensors and MNE’s fsaverage coordinates.

#### Inverse solution

Inverse solutions were obtained separately for Repeat and Novel epochs. All epochs were first baseline-corrected by subtracting the mean channel data of the baseline period (t = -0.5 s to t = 0 s from stimulus presentation) from all channels over the entire epoch duration, epoch-by-epoch. Each participant’s inverse operator was set up using a Minimum Norm Estimate (MNE) model given the forward model and with source orientation constrained to be orthogonal to the cortical mantle (*mne.minimum norm.make inverse operator*, setting the argument “fixed”=True). Next, inverse solutions (or source estimates) were calculated for each epoch. This required computation of a noise covariance matrix, which was calculated from the baseline segments of all epochs using *mne.compute covariance* with the argument “keep sample mean”=False. The function *mne.SourceEstimate.estimate_snr* was used to estimate the regularization parameter “lambda” for each participant in the overt naming experiment, averaging the peak estimates across participants. The inverse operator and noise covariance matrix served as inputs to *mne.minimum norm.apply inverse epochs* (with the argument “method”=dSPM and the estimated “lambda” =0.0004) to obtain induced source estimates for all epochs and all participants. The argument “pick_ori” was set to “None” so that the source estimates were signed.

### ROI source time series, and time-frequency (t-f) power

ROIs defined in fMRI data, which used SUMA’s std.141 surface mesh, were registered to the fsaverage surface. To co-register our source estimates defined on the participant’s anatomically constrained mesh and the fMRI masks defined on fsaverage, we used *mne.compute source morph*. Next, for each ROI, we extracted vertex-averaged induced time series by combining the time series of all of their sources using *mne.extract label time course* with argument mode set to “pca flip”. The same function/arguments were used to extract time series from the evoked source estimates. Time-frequency power spectra were obtained from each ROI timeseries using Morlet wavelets (*mne.time frequency.tfr array morlet*). We used frequencies 0.5 Hz and 1…100Hz separated by 1Hz intervals (via the argument “freqs”). We set the number of cycles of each wavelet (argument “n cycles”) to be a third of its frequency (e.g. for 30 Hz, 10 cycles were used) and arguments “use fft”=True and “output”=power. Finally, we *z*-scored the t-f power spectra per frequency for each epoch with respect to the baseline period (t = -0.5 s to t = 0 s). Note that these were calculated for both “induced” (one t-f power spectrum per ROI per epoch after first removing the ERP from each epoch) and “evoked” sources (a single evoked power time spectrum per ROI calculated on the ERP). To calculate average ROI induced power spectra for each participant, we averaged the power spectra over all epochs. In the end, this resulted in one induced and one evoked power spectrum for each ROI, per condition (Novel, Repeat), per participant. The process described above was the same for both overt and covert EEG data. It was also the same used to generate t-f power series of Repeat minus Novel differences projected for each vertex on the cortical surface as Supplementary Movies 1-4. Banded power estimates as a function of time for each condition, ROI, and participant were formed by further averaging the average ROI t-f power spectrum in each condition over the frequencies defining the bands (delta = 0.5-3 Hz, theta = 4-7 Hz, alpha = 8-12 Hz, beta = 13-29 Hz, gamma = 30-80 Hz).

### Region of interest (ROI) power analyses and correlations with fMRI RS and priming

Statistical comparisons across participants utilized paired *t*-tests for each ROI of Repeat- Novel power at each frequency and time sample. Comparisons per frequency band were then corrected for multiple comparisons using FDR (*q* < .05). Contiguous periods of “early” (Repeat > Novel) and “late” (Novel > Repeat) power differences in each frequency band that survived FDR correction were then defined as any period during the overall group effect lasting at least 50 ms for which the direction of the effect was the same. These periods were then averaged for each participant for subsequent correlations with fMRI repetition suppression and repetition priming effects (one “early” and one “late” average power difference for each participant and frequency band). Correlations (Spearman rank-order) were then carried out across participants between early and late power differences (Repeat-Novel) in each frequency band and ROI with fMRI repetition suppression (Repeat-Novel beta coefficients) estimated for the same ROI and the magnitude of repetition priming (Cohen’s *d* values). Multiple comparisons across early/late periods, frequency bands, and ROIs were corrected using FDR (*q* < .05). Spearman rank-order correlations (and partial correlations for follow-up analyses) were utilized to guard against any potential effects of strong leverage (outlying) data points or possible violations of normality. For replication purposes, Covert Naming power analyses utilized the time windows of significance from Overt Naming for the same ROIs (earliest time point to latest time point across frequencies), performing further FDR correction (*q* < .05) across time points within these windows.

## ACKNOWLEDGEMENTS

The views expressed in this article do not necessarily represent the view of the NIH, the Department of Health and Human Services, or the United States Government. We acknowledge funding support from the National Institute of Mental Health, NIH, Division of Intramural Research grant ZIAMH002920 (to AM). The funders had no role in study design, data collection and analysis, decision to publish, or preparation of the manuscript.

## SUPPLEMENTARY FIGURES AND TABLES

**Supplementary Fig. 1.**
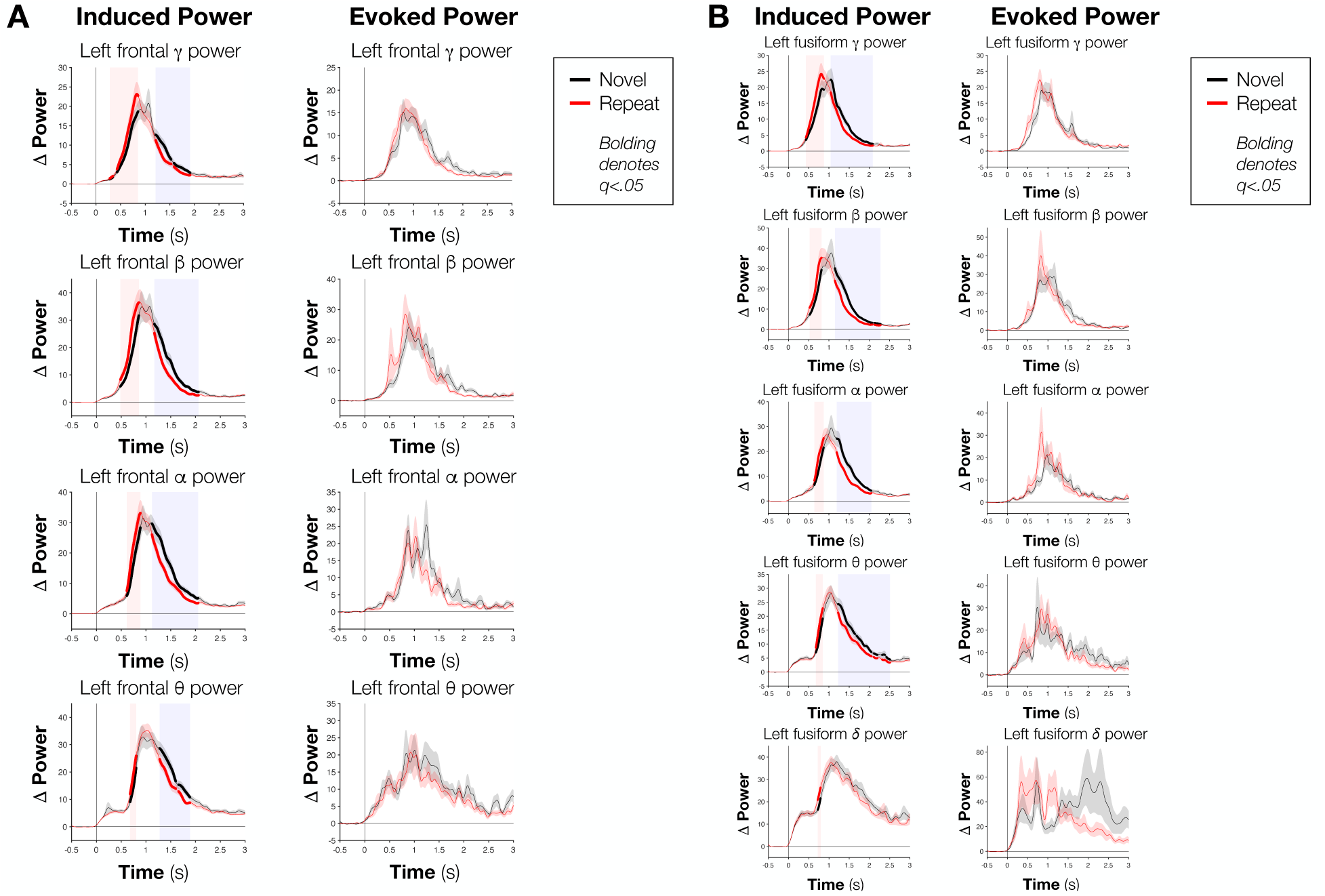
Effects of repeated object identification on induced versus evoked power for the fMRI-defined A) left frontal and B) left fusiform regions. Evoked power corresponds to power estimates after first averaging across trials in the time domain per condition, whereas induced power is calculated per trial after first subtracting the trial-averaged evoked response and then averaging across trials. FDR-corrected periods are shown with bolded lines, with red lines representing change in power relative to the baseline period for the Repeat condition and black lines representing the Novel condition.

**Supplementary Fig. 2.**
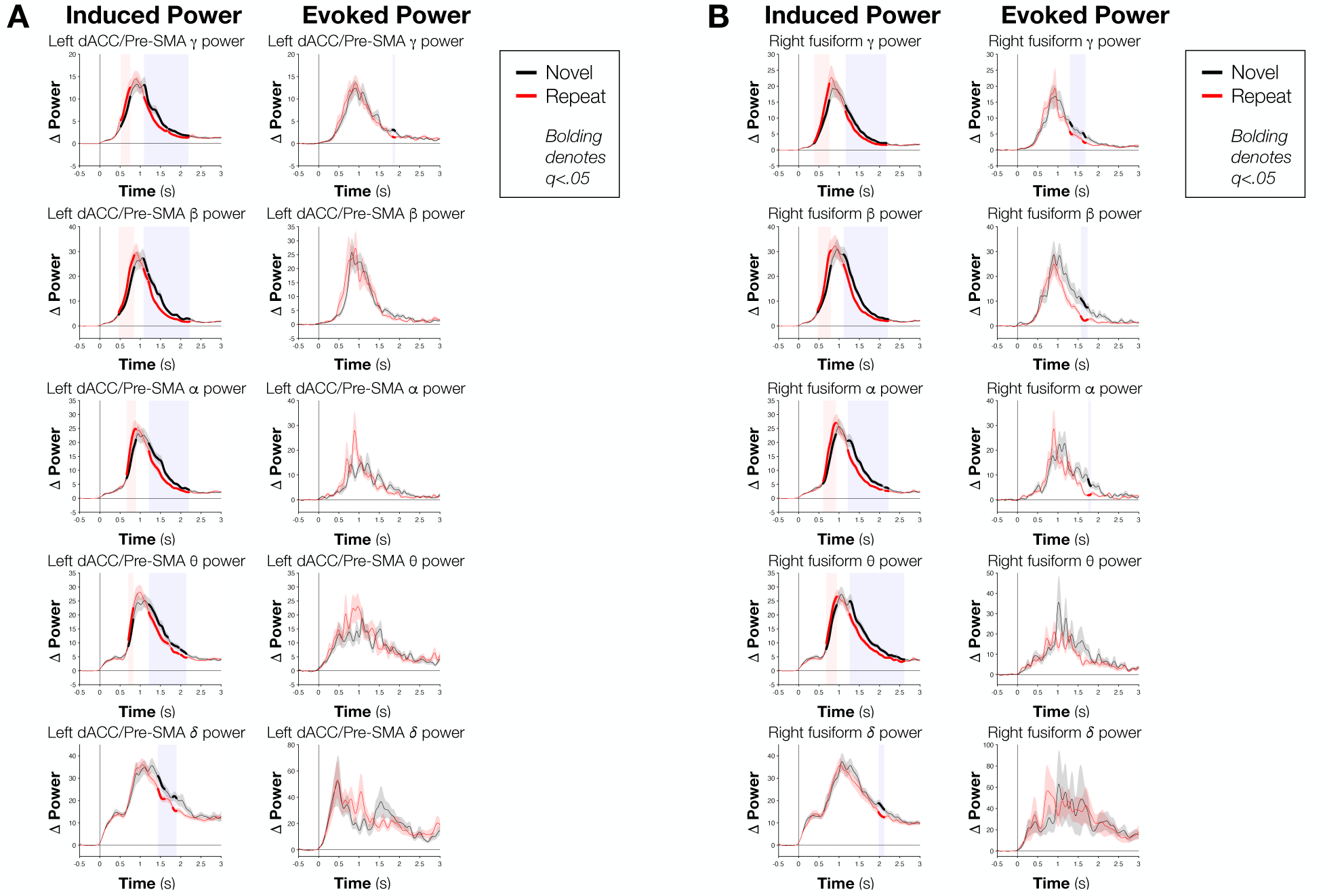
Effects of repeated object identification on induced versus evoked power for the fMRI-defined A) dorsal anterior cingulate/pre-supplementary motor area (dACC/Pre-SMA) and B) right fusiform regions. Evoked power corresponds to power estimates after first averaging across trials in the time domain per condition, whereas induced power is calculated per trial after first subtracting the trial-averaged evoked response and then averaging across trials. FDR-corrected periods are shown with bolded lines, with red lines representing change in power relative to the baseline period for the Repeat condition and black lines representing the Novel condition.

**Supplementary Fig. 3.**
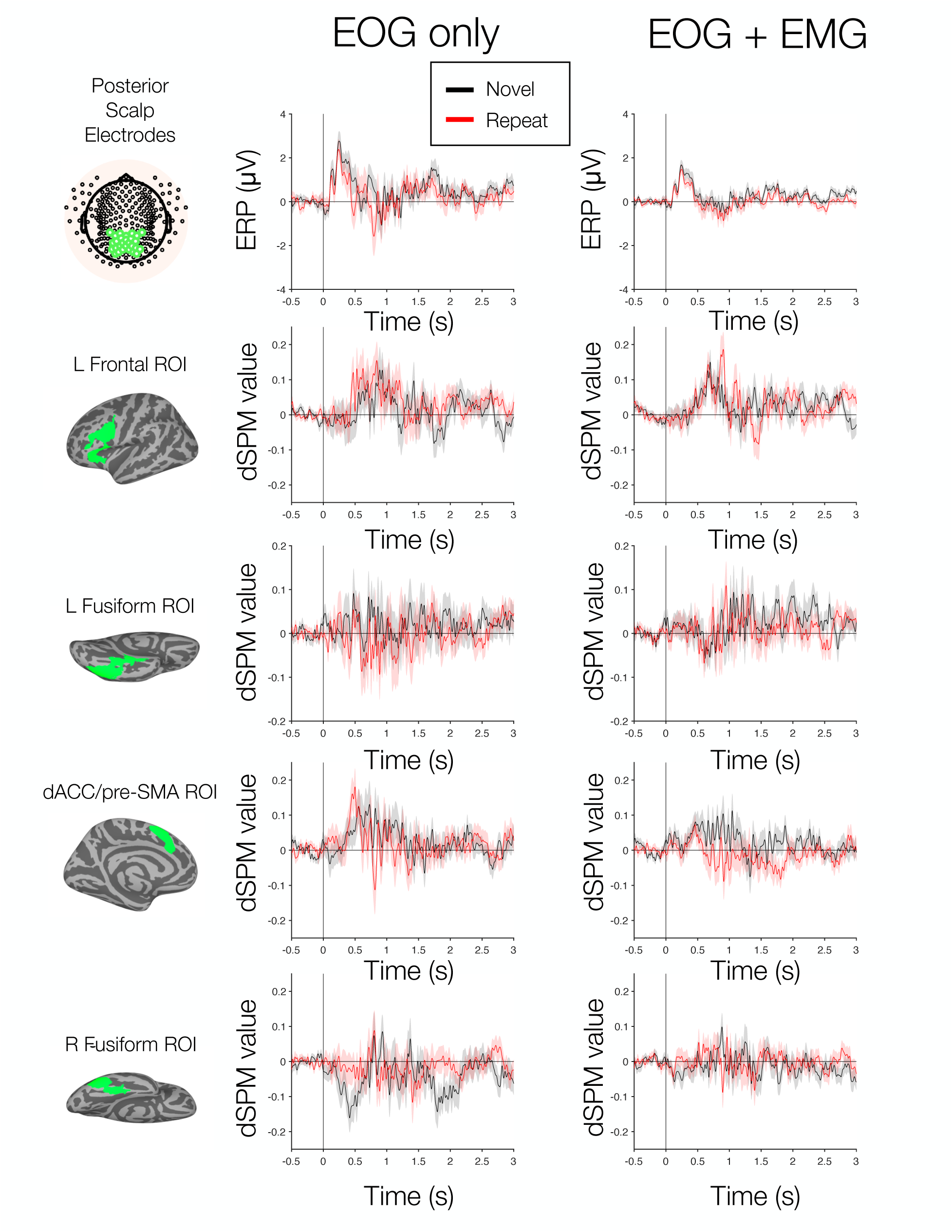
Event related potentials for EOG only (pipeline 1) versus EOG plus speech EMG pre-processing (pipeline 2). Novel and Repeat condition ERPs (black and red lines, respectively) are shown in the top row for the selection of posterior scalp electrodes (graphic on left), then for the source estimated time series of the fMRI-defined regions (left frontal, left fusiform, left dACC/pre-SMA, and right fusiform regions). The amplitudes of the ERPs are reduced near the time of the average response time (∼ .9 seconds) and .5 seconds before for the EOG+EMG pre-processing relative to the EOG only pre-processing. The estimated ERPs are unitless for the fMRI-defined regions due to the normalization by noise covariance periods.

**Supplementary Fig. 4.**
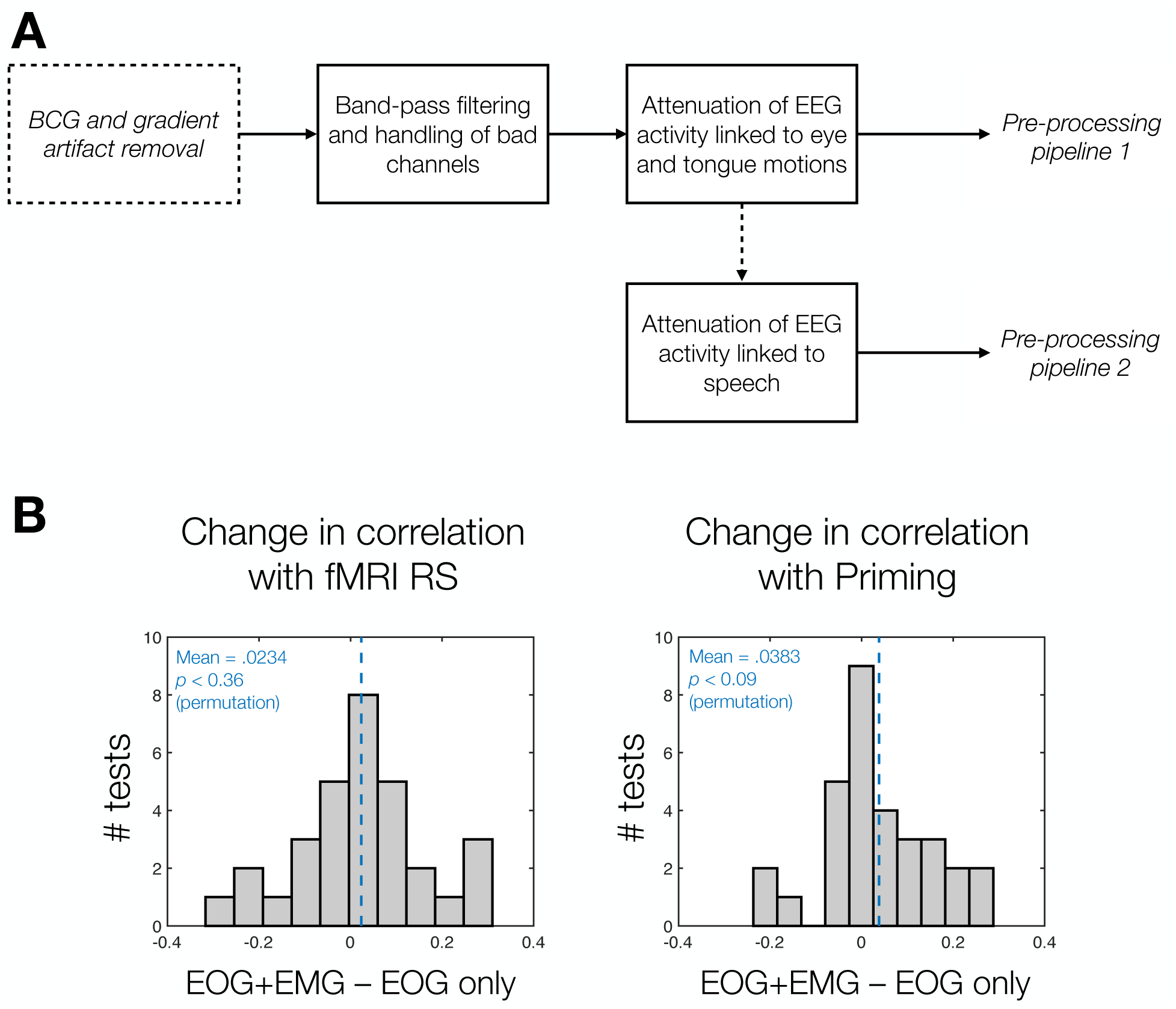
Schematic of different pre-processing pipeline and how these impact correlations of induced power differences (Repeat - Novel) for the early and late periods with fMRI RS and priming. A) Pipeline 1 (“EOG only”) regressed signals from electrodes near the eyes and upper jaw to remove EOG and tongue EMG signals, whereas Pipeline 2 (“EOG+EMG”) additionally removed signals from the scalp electrodes based on ICAs calculated near the time of the speech response (-0.2 to +0.2 s relative to the RT). See Materials and Methods: EEG Preprocessing for further details. B) Correlations of induced power differences (Repeat - Novel) for the early and late periods with fMRI RS and priming show little change or slightly larger correlations in the EOG+EMG preprocessing (pipeline 2) relative to the EOG only preprocessing (pipeline 1). Shown are frequency histograms of the change in correlation between pipelines across all tests, with mean change values near zero for correlations with fMRI RS and marginally positive for correlations with priming (*p* < .09, permutation test). Negative correlations were first rectified to positive values using absolute value for the purposes of these calculations (e.g. positive values in the histogram reflect a stronger correlation – either originally positive or negative – in the EOG+EMG relative to the EOG only preprocessing). Permutation testing (10,000 iterations) was conducted by randomly flipping the signs of the correlation differences and recalculating the mean differences.

**Supplementary Fig. 5.**
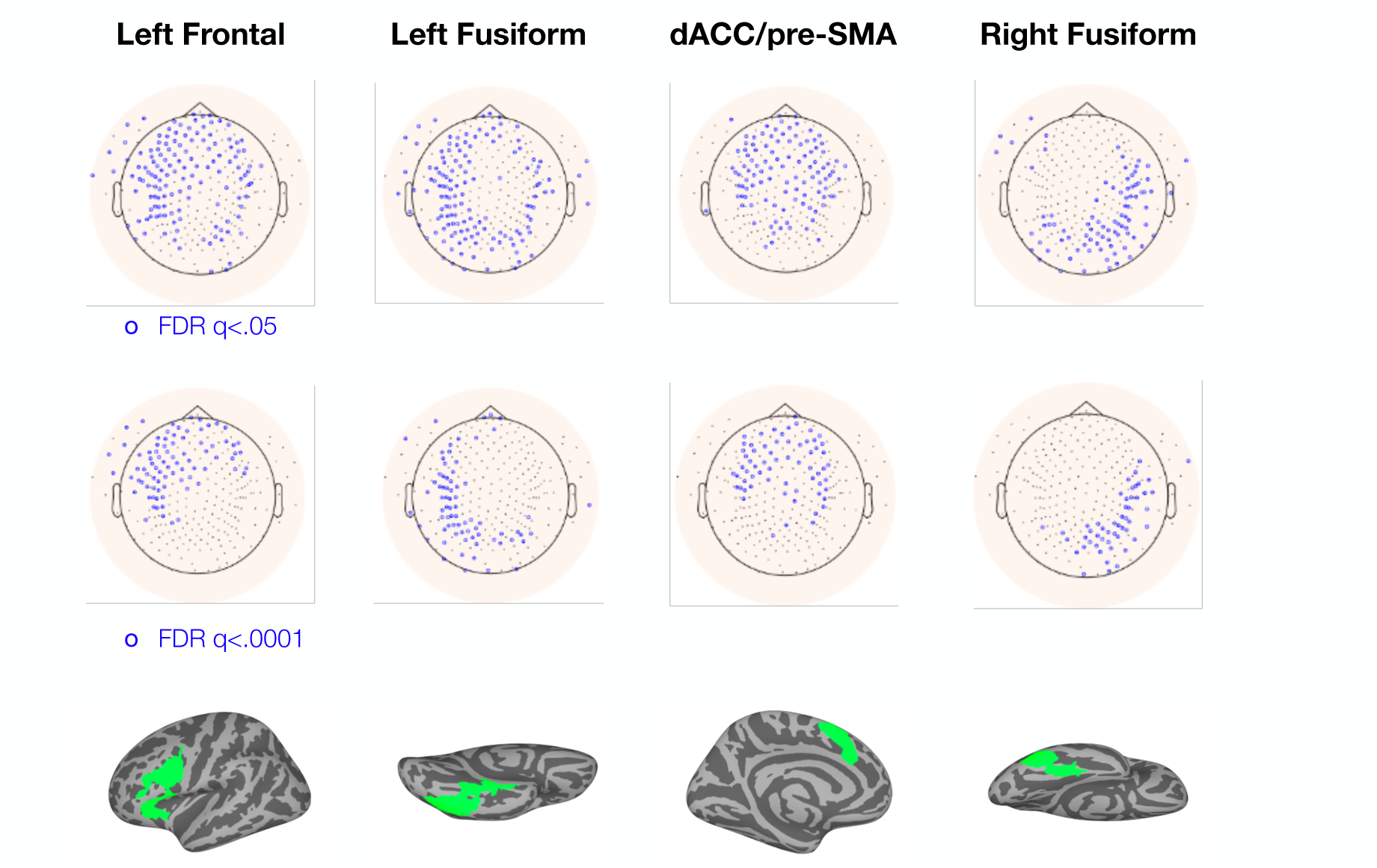
Mapping source regions to their most heavily weighted scalp electrodes. Shown are two FDR-corrected significance levels: FDR *q* < .05 (minimum thresholding for significance) and *q* < .0001 (more stringent thresholding). Some degree of spatial selectivity is observed under both thresholds, although even at the more stringent threshold (*q* < .0001) there is overlap of the identified scalp electrodes across regions – highlighting the more limited spatial resolution of EEG. Electrodes were identified by taking the electrode weights associated with each source vertex in the source estimation process (see Materials and Methods), normalizing them by the standard deviation across those weights, and then finding the mean weight with each surface electrode for all the vertices belonging to a given fMRI-defined region. One-sample *t*- tests across participants were then carried out for each scalp electrode per fMRI-defined region. Multiple comparisons were corrected by FDR. ERPs and induced power differences for these electrode selections are shown in Supplementary Figs. 6-9. While induced power differences appear similar across frequencies and fMRI-defined regions, the ERPs for these electrode selections have distinctive shapes, consistent with a moderate degree of spatial selectivity of the scalp EEG signals.

**Supplementary Fig. 6.**
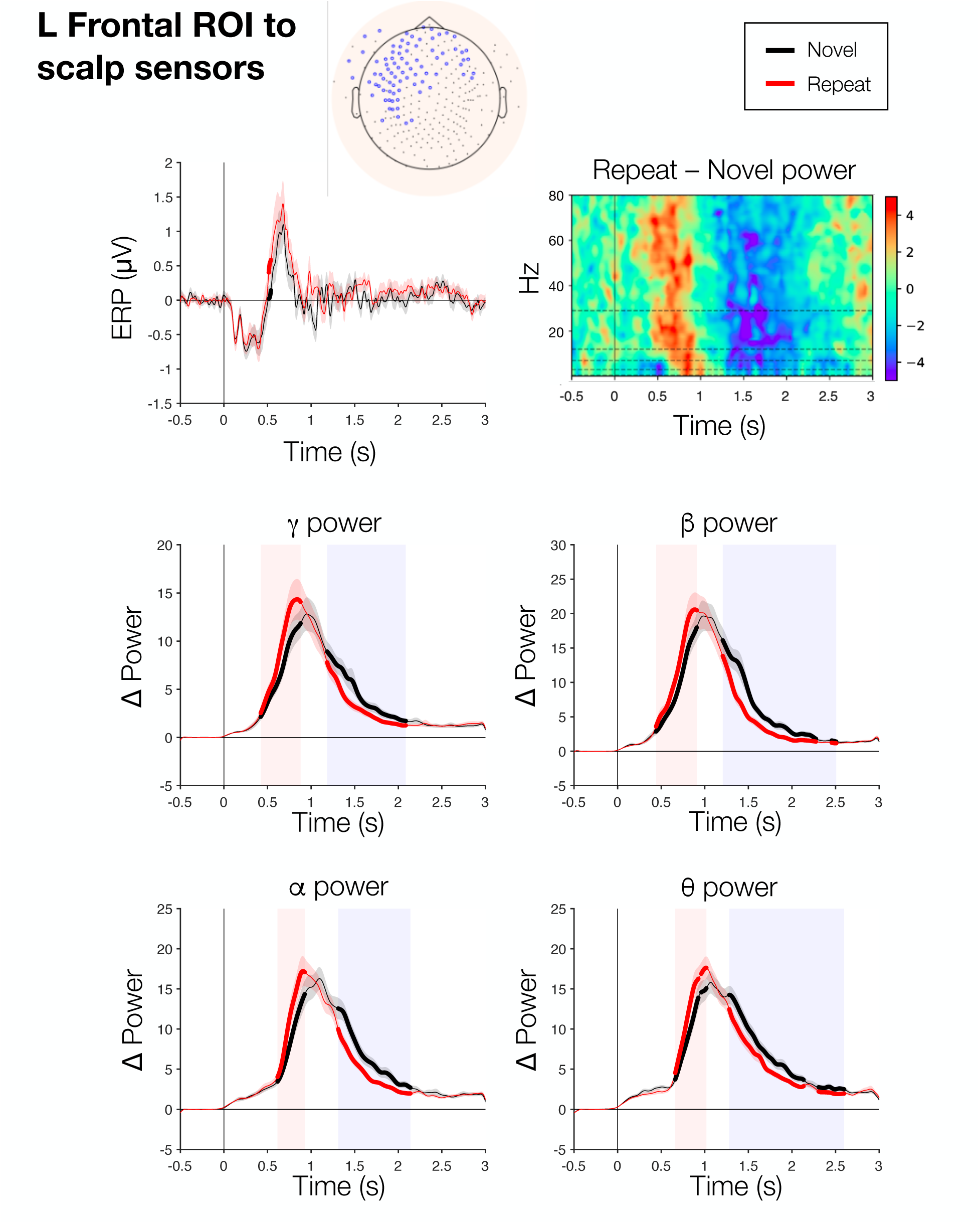
Companion scalp electrode analyses for the fMRI-defined left frontal region (FDR *q* < .0001 from Supplementary Fig. 5). Average ERPs for the defined collection of electrodes (see inset at top) for the Novel (black) and Repeat (red) conditions. Repeat-Novel time by frequency induced power differences are shown in the upper right (paired *t*-tests across participants), highlighting the same “early” (Repeat > Novel) and “late” (Novel > Repeat) periods as in the source-estimated results in Fig. 2. Banded power analyses (Novel vs. Repeat) are then shown below for the gamma (30-80 Hz), beta (13-29 Hz), alpha (8-12 Hz) and theta (4-7 Hz) frequency bands, with FDR-corrected periods of significance indicated with bolded lines. While no FDR-corrected periods were detected in the ERPs, robust induced power differences are observed across frequencies.

**Supplementary Fig. 7.**
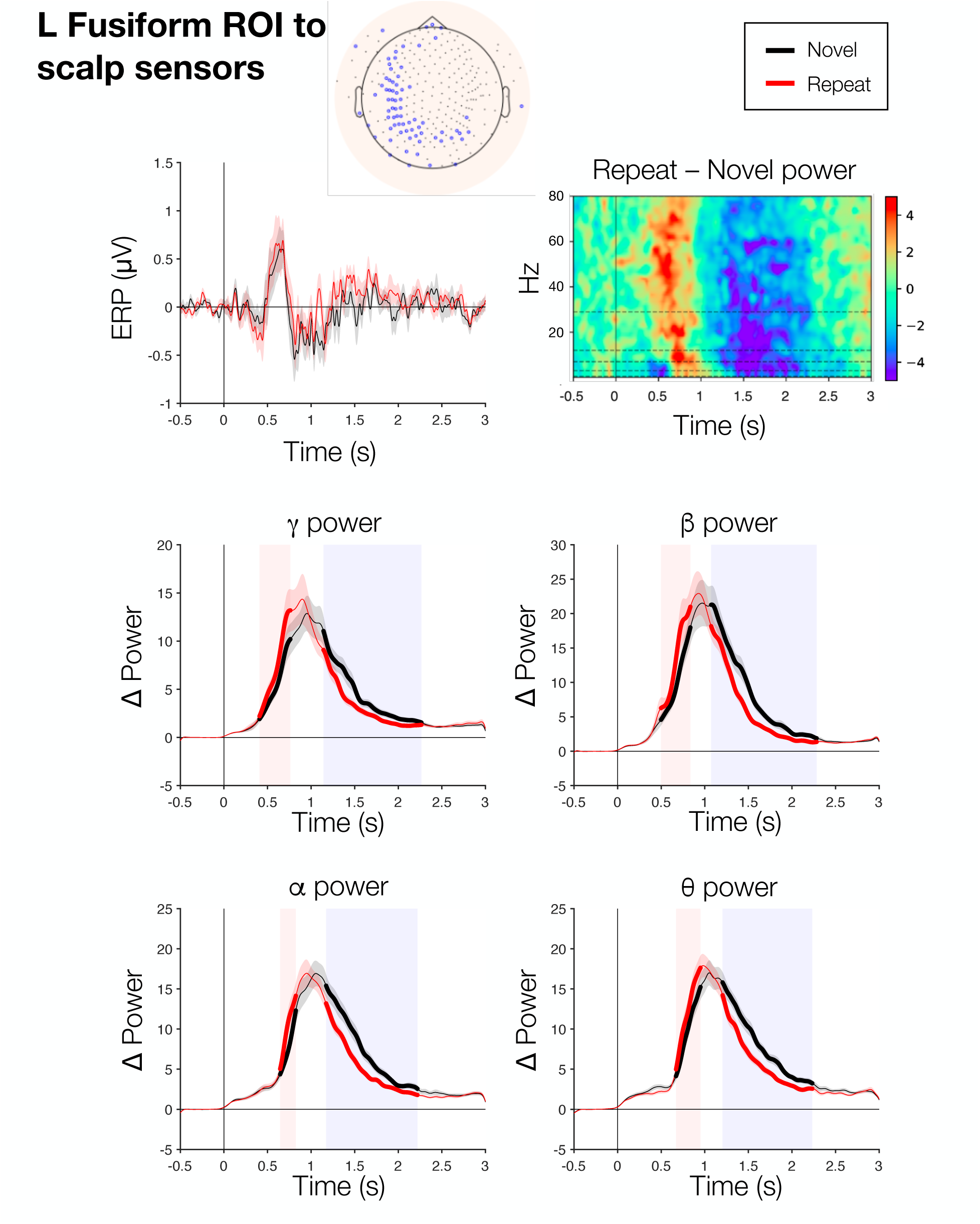
Companion scalp electrode analyses for the fMRI-defined left fusiform region (FDR *q* < .0001 from Supplementary Fig. 5). Average ERPs for the defined collection of electrodes (see inset at top) for the Novel (black) and Repeat (red) conditions. Repeat-Novel time by frequency induced power differences are shown in the upper right (paired *t*-tests across participants), highlighting the same “early” (Repeat>Novel) and “late” (Novel>Repeat) periods as in the source-estimated results in Fig. 2. Banded power analyses (Novel vs Repeat) are then shown below for the gamma (30-80 Hz), beta (13-29 Hz), alpha (8-12 Hz) and theta (4-7 Hz) frequency bands, with FDR-corrected periods of significance indicated with bolded lines. While no FDR-corrected periods were detected in the ERPs, robust induced power differences are observed across frequencies.

**Supplementary Fig. 8.**
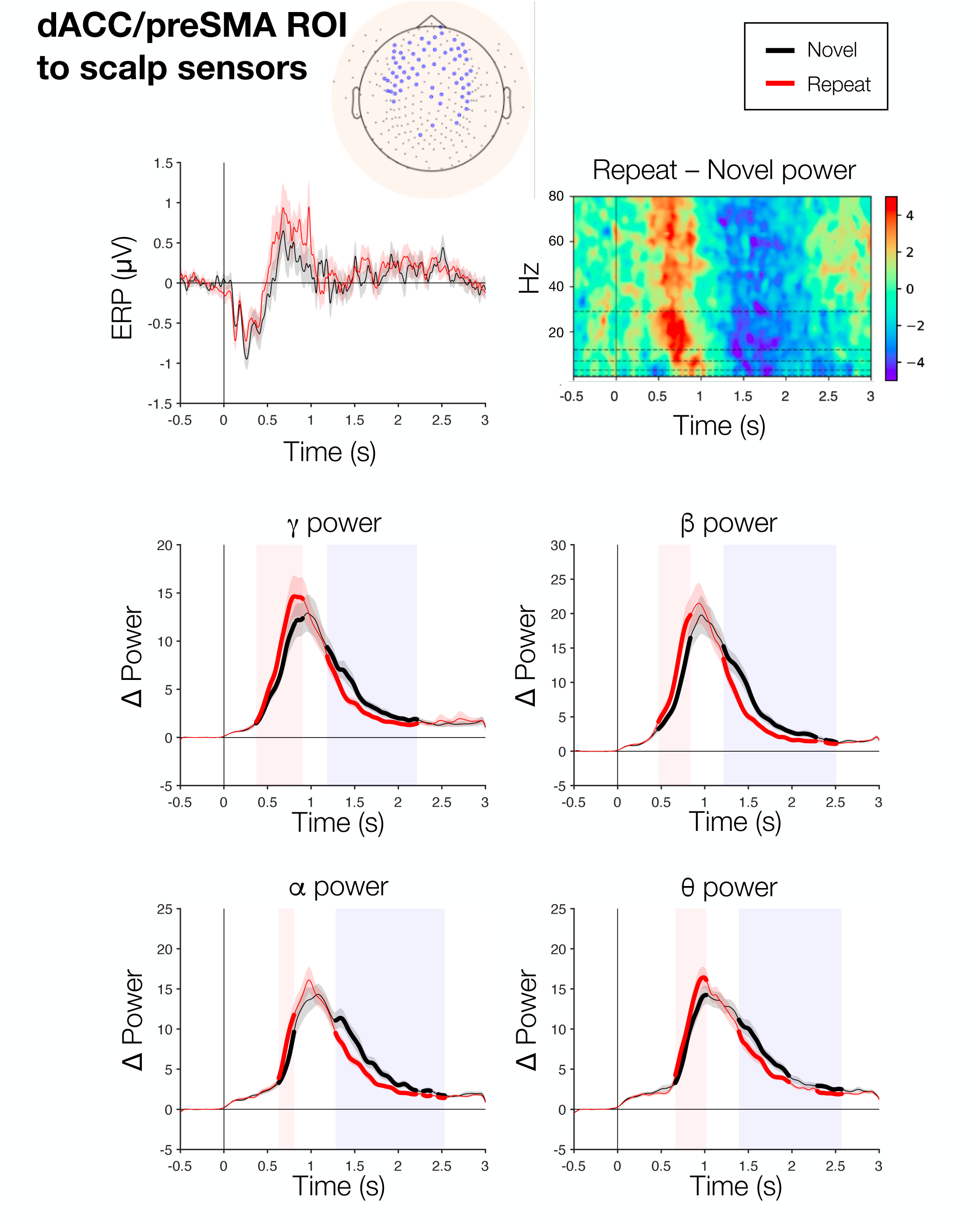
Companion scalp electrode analyses for the fMRI-defined left dACC/pre-SMA region (FDR *q* < .0001 from Supplementary Fig. 5). Average ERPs for the defined collection of electrodes (see inset at top) for the Novel (black) and Repeat (red) conditions. Repeat-Novel time by frequency induced power differences are shown in the upper right (paired *t*-tests across participants), highlighting the same “early” (Repeat>Novel) and “late” (Novel>Repeat) periods as in the source-estimated results in Fig. 2. Banded power analyses (Novel vs Repeat) are then shown below for the gamma (30-80 Hz), beta (13-29 Hz), alpha (8-12 Hz) and theta (4-7 Hz) frequency bands, with FDR-corrected periods of significance indicated with bolded lines. While no FDR-corrected periods were detected in the ERPs, robust induced power differences are observed across frequencies.

**Supplementary Fig. 9.**
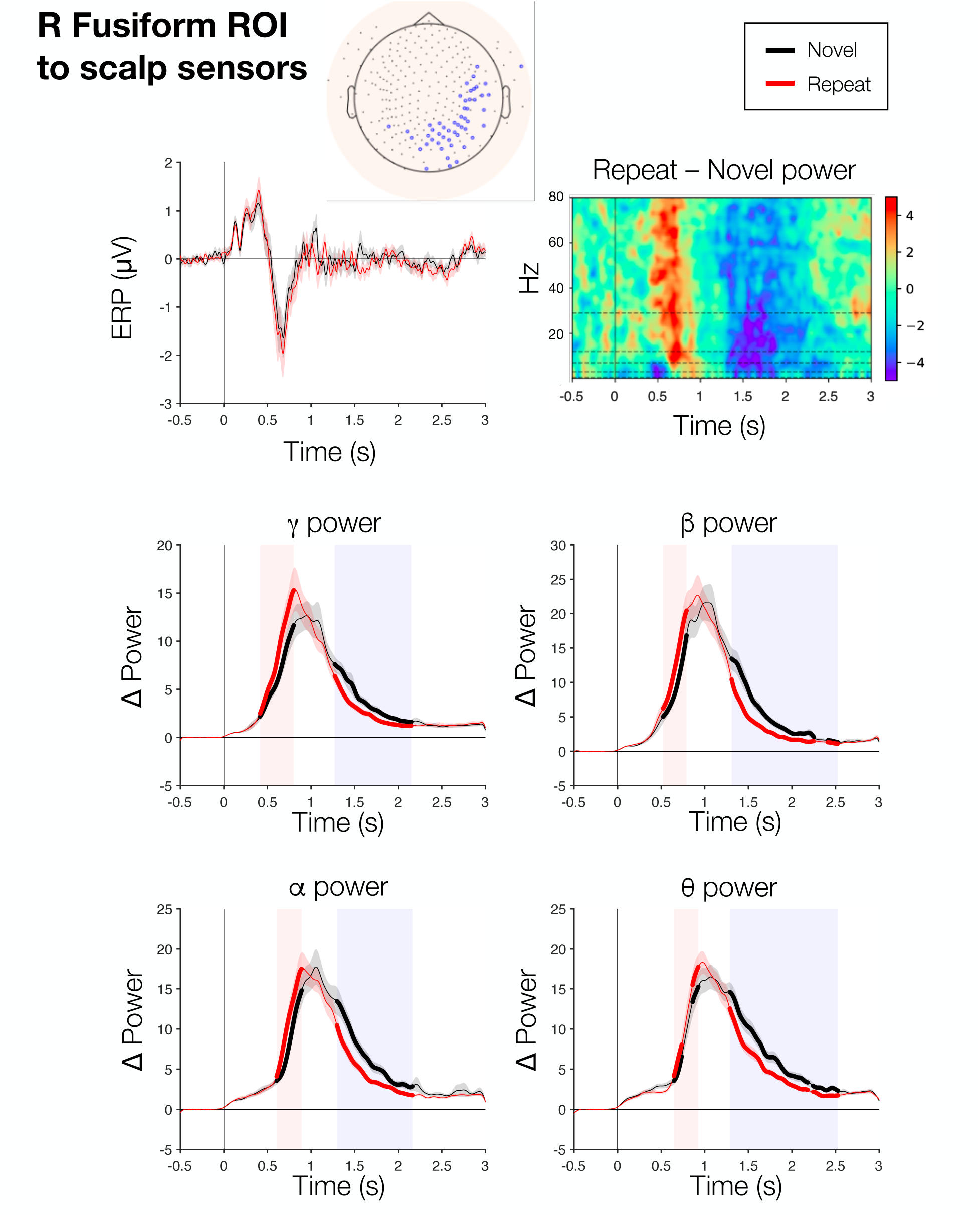
Companion scalp electrode analyses for the fMRI-defined right fusiform region (FDR *q* <.0001 from Supplementary Fig. 5). Average ERPs for the defined collection of electrodes (see inset at top) for the Novel (black) and Repeat (red) conditions. Repeat-Novel time by frequency induced power differences are shown in the upper right (paired *t*-tests across participants), highlighting the same “early” (Repeat>Novel) and “late” (Novel>Repeat) periods as in the source-estimated results in Fig. 2. Banded power analyses (Novel vs Repeat) are then shown below for the gamma (30-80 Hz), beta (13-29 Hz), alpha (8-12 Hz) and theta (4-7 Hz) frequency bands, with FDR-corrected periods of significance indicated with bolded lines. While no FDR-corrected periods were detected in the ERPs, robust induced power differences are observed across frequencies.

**Supplementary Fig. 10.**
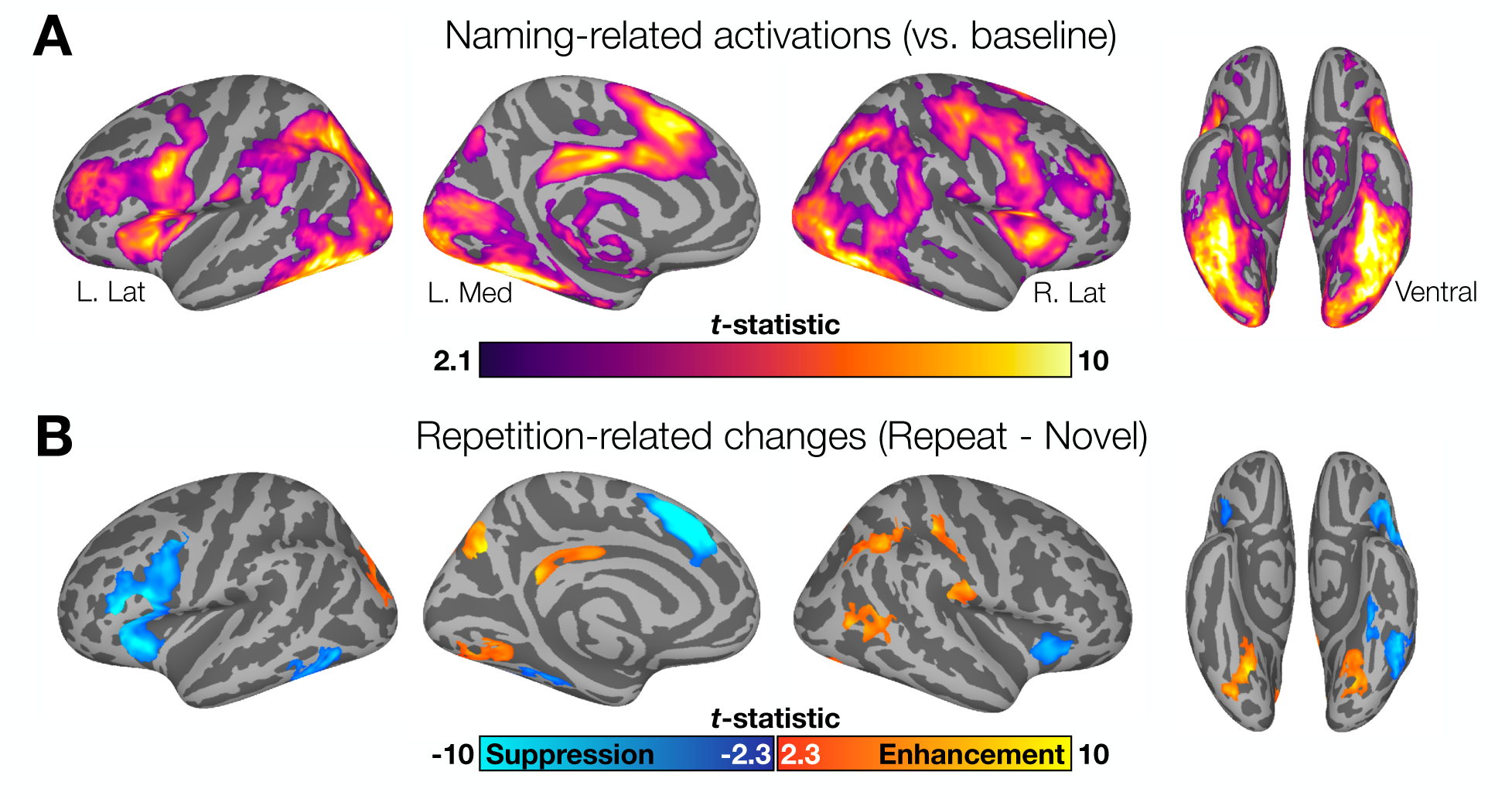
Vertexwise results of Naming-related activations and Repetition effects in the covert naming experiment. (A) fMRI identified significant activations in multiple regions across the cortex, consistent with those observed in the overt naming task. (B) Within Naming- activated regions, significant repetition suppression was noted in frontal and fusiform cortex and were generally left-lateralized. Effects are somewhat attenuated when compared to those in the Overt data, potentially due a combination of the covert nature of the task and the smaller sample size. For display purposes, the map of repetition effects is set to *q* < .05, inclusively masked by regions identified in the earlier Naming-related activation map, while maintaining the same cluster size minimum.

